# CryoFIB milling large tissue samples for cryo-electron tomography

**DOI:** 10.1101/2022.10.03.510728

**Authors:** Sihan Wang, Heng Zhou, Wei Chen, Yifeng Jiang, Xuzhen Yan, Hong You, Xueming Li

## Abstract

Cryo-electron tomography (cryoET), a powerful tool for exploring the molecular structure of large organisms. However, technical challenges still limit cryoET applications on large samples. In particular, locating and cutting out objects of interest from a large tissue sample is an important but difficult step. In this study, we report a sample thinning strategy and workflow for tissue samples based on cryo-focused ion beam (cryoFIB) milling. This workflow provides a full solution for isolating objects of interest by starting from a millimeter-sized tissue sample and ending with hundred-nanometer thin lamellae. The workflow involves sample fixation, pre-sectioning, a two-step milling strategy, and locating the object of interest using cellular secondary electron imaging (CSEI). The two-step milling strategy introduces a coarse milling method to solve the milling efficiency problem for samples as thick as tens of microns, followed by a fine milling method to create a furrow-ridge structure. The furrow-ridge structure guarantees the generation of large, thin lamellae with enhanced mechanical stability and charge-reducing design. CSEI is highlighted in the workflow, which provides conventional, on-the-fly locating during cryoFIB milling. Tests of the complete workflow were conducted to demonstrate the high efficiency and high feasibility of the proposed method.

## Introduction

Cryo-electron tomography (cryoET) is a rapidly developing and popular technique that enables the direct study of high-resolution structures of biological macromolecules and their interactions in cells and tissues *in situ*^1, 2^. However, due to the strong interactions between electrons and biological material, frozen hydrated biological samples must be sliced into thin lamellae, typically not exceeding 200–300 nm thick^3^, ahead of cryoET imaging. Finding and cutting out a thin lamella containing the objects of interest from a millimeter-size tissue sample is one of the major challenges of cryoET, and effectively limits the application of this technology on large tissue samples.

Cryo-focused ion beam (cryoFIB) milling^4, 5^ is a promising technique for cryoET sample preparation. In contrast to conventional cryo-ultramicrotomy using a diamond knife^6^, cryoFIB can avoid mechanical damage to the sample during the sectioning process. However, the ion beam has very low efficiency in removing large volumes. Consequently, cryoFIB is mostly used for samples with an initial thickness of several micrometers^2^. To improve the efficiency and enable cryoFIB use on large tissue samples, attempts have been made to combine cryo-ultramicrotomy with cryoFIB. For example, Hayles et al. froze cells in a copper tube using high-pressure freezing (HPF), then trimmed the tube using cryo-ultramicrotomy to a specific shape with suitable thickness for cryoFIB milling^7^. Zhang et al. froze a large tissue sample in a special cryo-carrier and subsequently trimmed the cryo-carrier together with the sample using cryo-ultramicrotomy to achieve an initial thickness of 20 μm for further cryoFIB milling^8^. A technique called cryo-lift out has been used to extract a small volume from a large sample with the assistance of cryoFIB and then milled using the conventional cryoFIB method^9^. While these methods can be applied to large tissue samples, the complicated operational procedures required are a major disadvantage as only well-trained users can successfully apply these methods, and total processing times are counted in days.

Another key issue involves locating objects of interest before or during the cryoFIB process. The field of view of cryoET is limited within a thin lamella with micrometer width. Locating and estimating the accurate three-dimensional (3D) position of such a small volume within a large bulky sample is required, but is difficult to accomplish during milling. Cryo-correlative light and electron microscopy (cryoCLEM)^10, 11^ is a popular solution for this problem that uses sophisticated fluorescence microscopes, primarily confocal and super-resolution fluorescence microscopes, to assist cryoFIB milling and cryoET analysis. In most applications to date, cryoCLEM has been performed outside the cryoFIB instrument; hence, cryoCLEM has been unable to achieve on-the-fly locating. Attempts to enable on-the-fly locating by integrating fluorescence microscopy into the cryoFIB instrument^12^ have been significantly limited by poor fluorescence imaging resolution due to the long working distance of the optical lens and limited installation space inside the cryoFIB instrument. In reports of cryoFIB-SEM block-face imaging^13, 14, 15^ and our accompanying work^16^, cellular secondary electron imaging (CSEI) allows direct visualization of the ultrastructure of cells. Therefore, we hypothesized that CSEI has the potential to enable on-the-fly locating and assist cryoFIB milling for large tissue samples.

In this study, we report a cryoFIB milling method and workflow for frozen hydrated tissue samples. This method allows for high-efficiency milling of large frozen hydrated tissue samples with on-the-fly locating of objects of interest, without the need for extra locating equipment. This workflow covers all sample preparation steps, including fixation, pre-sectioning, CSEI locating, and coarse and fine milling. Additionally, the charging issue commonly observed in large tissue lamella is remarkably weakened by virtue of a furrow-ridge structure prepared in the fine milling step. Finally, an example of the complete workflow is illustrated by targeting collagen fibrils in mouse liver tissue.

## Results

### Overall workflow for high-efficiency milling of large tissue samples

The initial size of tissue samples is often >1 mm, whereas the workable sample size for HPF and cryoFIB is much smaller. The size requirement gap among these processing steps and size-reducing efficiency should be carefully considered. The optimal thickness for HPF must be ≤200 μm^17^ to ensure that the sample is fully vitrified. The workable thickness of cryoFIB determines the milling time, which increases significantly with the initial sample thickness; generally, 1–2 hours are required for samples of 1–5 μm thick, and several hours or even days may be required for multicellular organisms that are tens of microns thick. In addition, the ion beam current is a key factor that determines the milling speed. For example, if cutting a window on a sample takes a few minutes using nA-level current, many hours would be required using pA-level current. Combining these analyses, we designed a workflow to achieve high-efficiency milling (**Fig. 1**).

**Figure 1.**
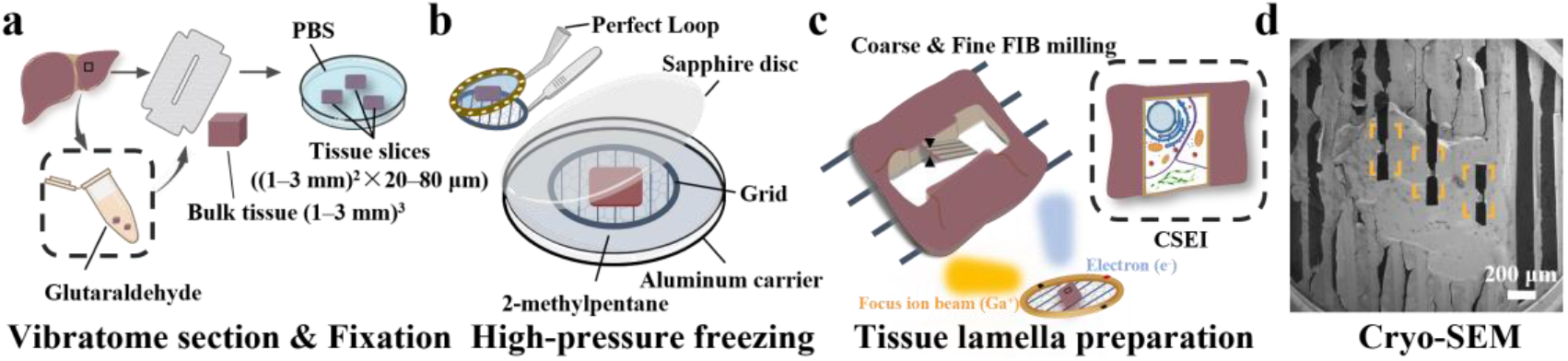
Schematic illustration of the overall workflow. **a**, Chemical fixation and pre-sectioning starting from a large tissue. **b**, High-pressure freezing (HPF) on a grid with parallel bars using 2-methylpentane. A Perfect Loop (Diatome, DZ8) is used to transfer the sliced sample to the grid. **c**, CryoFIB milling with CSEI-based locating. The incident FIB beam should be parallel to the grid bars. **d**, Typical SEM image of the sample with multiple lamellae (yellow boxes) prepared on one tissue block.

The first step was the pre-sectioning, which used a mechanical method to slice the initial sample to a workable thickness for HPF and cryoFIB milling (**Fig. 1a**). A cuboid with an edge length of 1-3 mm (smaller than a standard 3-mm grid) was cut out of the original bulky tissue and used as the initial sample for pre-sectioning. Chemical fixation, while optional, was often used ahead of pre-sectioning to minimize damage induced by the cellular contents from broken cells^18^(**Supplementary Fig. 1a and b**). Additionally, fixation may have increased the hardness of the tissue samples, thereby improving the mechanical sectioning accuracy (discussed later).

The second step involved loading the sample to a grid and performing HPF (**Fig. 1b**). 2-methylpentane, which sublimes at -150°C under high vacuum, was the preferred cryo-protectant for HPF as it can be easily removed by warming after HPF^19^. We recommend using a grid with parallel bars (**Fig. 1b**), which lacks grid bars in one direction, to avoid obstructing the ion and electron beam during cryoFIB milling and subsequent data collection of tilt series.

The third step involved multiple platinum (Pt) coatings and a two-step milling strategy composed of coarse and fine milling sub-steps (**Fig. 1c**). CSEI was used throughout the whole milling procedure to determine the milling positions. The Pt coating included an organometallic Pt deposition to protect the sample from ion beam radiation damage and two sputter coating to improve the surface electric conductivity (**Supplementary Protocol**). Coarse milling was performed to rapidly remove large sample volume using a large ion current at nA level, while fine milling was used to generate the final lamella with a special lamella structure (discussed later). With this milling strategy, the preparation of a tissue lamella as large as 20×50 μm typically required only 6 hours (**Supplementary Tables 1 and 2**). While the total milling time was longer than that for a thin sample, the larger lamella size contributed more cryoET tomograms per single lamella (**Supplementary Table 3**). The milling workflow also allowed for multiple positions to be milled together on the same grid (**Fig. 1d**).

### Chemical fixation and pre-sectioning

The pre-sectioning step was used to generate an initial workable thickness for HPF and cryoFIB milling. We recommend the use of a vibratome, which ensures the reproducibility of the sectioning process. The recommended section thickness was 20–80 μm; thinner sections require less time for subsequent cryoFIB milling.

Pre-sectioning often failed for very soft tissue samples; therefore, increasing the sample hardness by chemical fixation became necessary. We tested the success rate following different fixation times using 2.5% glutaraldehyde on mouse liver samples at room temperature (**Supplementary Fig. 1c**). Longer fixation times allowed more glutaraldehyde to diffuse into the sample, resulting in a less broken central region; in the samples with shorter fixation times, the central region was often largely broken or appeared brighter (**Supplementary Fig. 1c**). Note that even though the central region was broken, the surrounding region was often still large enough for cryoFIB milling. Therefore, a fixation time of 10–30 min was used in subsequent experiments.

Notably, we observed that the actual sample thickness was often larger than the expected thickness based on the vibratome settings used (**Supplementary Fig. 1d and e**). Additionally, the thickness was sometimes uneven in different areas of the same sample (**Supplementary Fig. 1f**). These factors should be considered when designing the experiments.

### On-the-fly locating using cellular secondary electron imaging

Secondary electron imaging is a basic function built into the cryoFIB instrument and is used to observe the surface topography and assist with cryoFIB milling. Under some settings, secondary electron imaging can capture cellular contrast on a flat surface created by cryoFIB milling. In our accompanying work^16^, we introduced CSEI to facilitate on-the-fly locating during cryoFIB milling. The imaging mechanism of CSEI is related to the secondary electron emission efficiency, the electrical conductivity of the bulk sample, and the surface charge under primary electron radiation. CSEI captures different contrast in cellular contents with different water content, membranes, and protein condensates. Importantly, CSEI can stably and rapidly display cellular features on the surface created by different ion beam currents.

The locating process with CSEI on tissue samples is simple and convenient, achieved by continuously imaging the fresh surface exposed by cryoFIB milling and targeting the objects of interest based on visible cellular features. We used mouse liver tissue as an example to test the locating workflow. First, we randomly selected a region (**Fig. 2a**) and milled a window of 80×20 μm under the cryoFIB view with a shallow beam angle of 18° and a strong ion beam current of 5 nA (**Fig. 2b**). On the generated surface, the ultrastructure of liver cells, such as membranes and various organelles, was clearly visible (**Fig. 2c**). Such imaging can be performed throughout the milling process.

**Figure 2.**
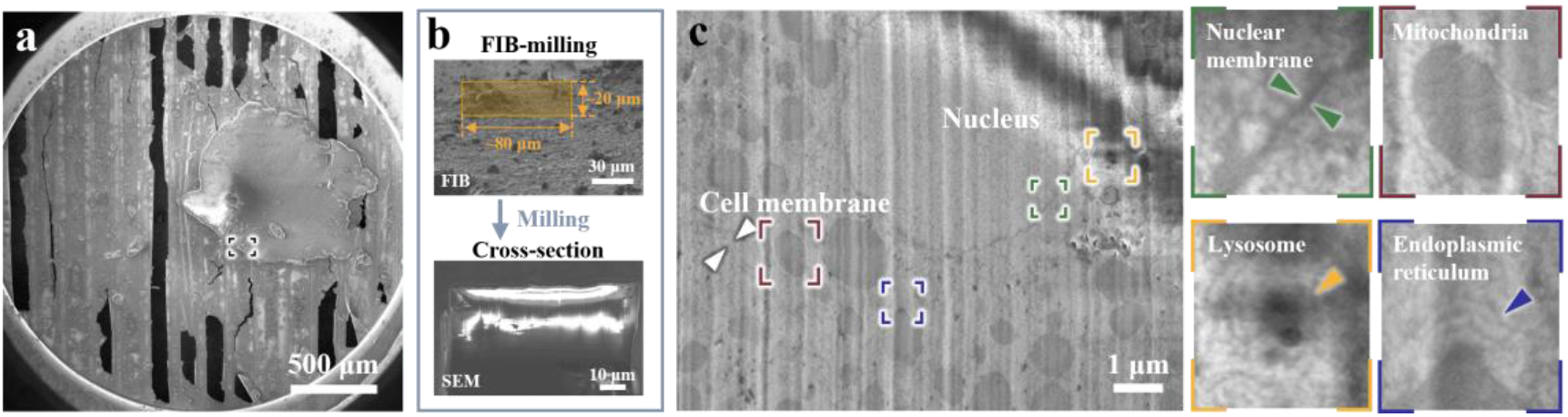
CSEI-based locating on a liver slice. **a**, Selected region (black box) for CSEI on a liver tissue slice, shown by low-magnification SEM. **b**, Large window (orange box in the top image, under FIB view) milled for CSEI (bottom image). **c**, Magnified CSEI area on the surface created by cryoFIB milling of (**b**) showed high contrast features of liver cells. The cell membrane (white arrow) and multiple organelles (colorful boxes and corresponding magnified images shown on the right) in liver cells were observed.

### Coarse milling to rapidly remove large volumes

The ion beam mills the sample along a small glancing angle nearly parallel to the sample surface; therefore, the volume to be removed increases significantly with increasing sample thickness. We considered two strategies to accelerate the milling: removing the volume around the region of interest and using high ion beam currents when possible. The bulky sample that remains around the final lamella forms a wall tens of microns high; this poses a problem for thick samples as it often shelters the electron beam during cryoET data collection.

We designed the coarse milling process with two sub-steps (see **Methods**). In the first sub-step, we milled the sample at a large incident angle (such as 48°), aiming to remove the volumes at the front and back ends of the region of interest (**Fig. 3a**); this sub-step effectively shortened the milling depth for the following milling step with a low ion beam current. The maximum ion beam current is recommended to maximize the milling speed. In our test (**Supplementary Fig. 2**), removing the volumes in two windows of 80×100 μm (under FIB view) on an ∼80 μm thick sample took less than 1 hour when using a beam current of 65 nA. In the second sub-step, we created a stepped edge around the lamella to avoid possible beam sheltering (**Fig. 3b and c**). In addition to the two sub-steps, one side of the lamella was disconnected from the bulky sample (**Fig. 3b, and c**). This was achieved by either completely removing the bulky sample on one side (**Supplementary Fig. 3**) or milling out a gap of 10 μm between the lamella and the remaining bulky sample (**Supplementary Fig. 4**). This step was essential to avoid crashing between the lamella and the remained sample in the future sample transfer (**Supplementary Fig. 5**).

**Figure 3.**
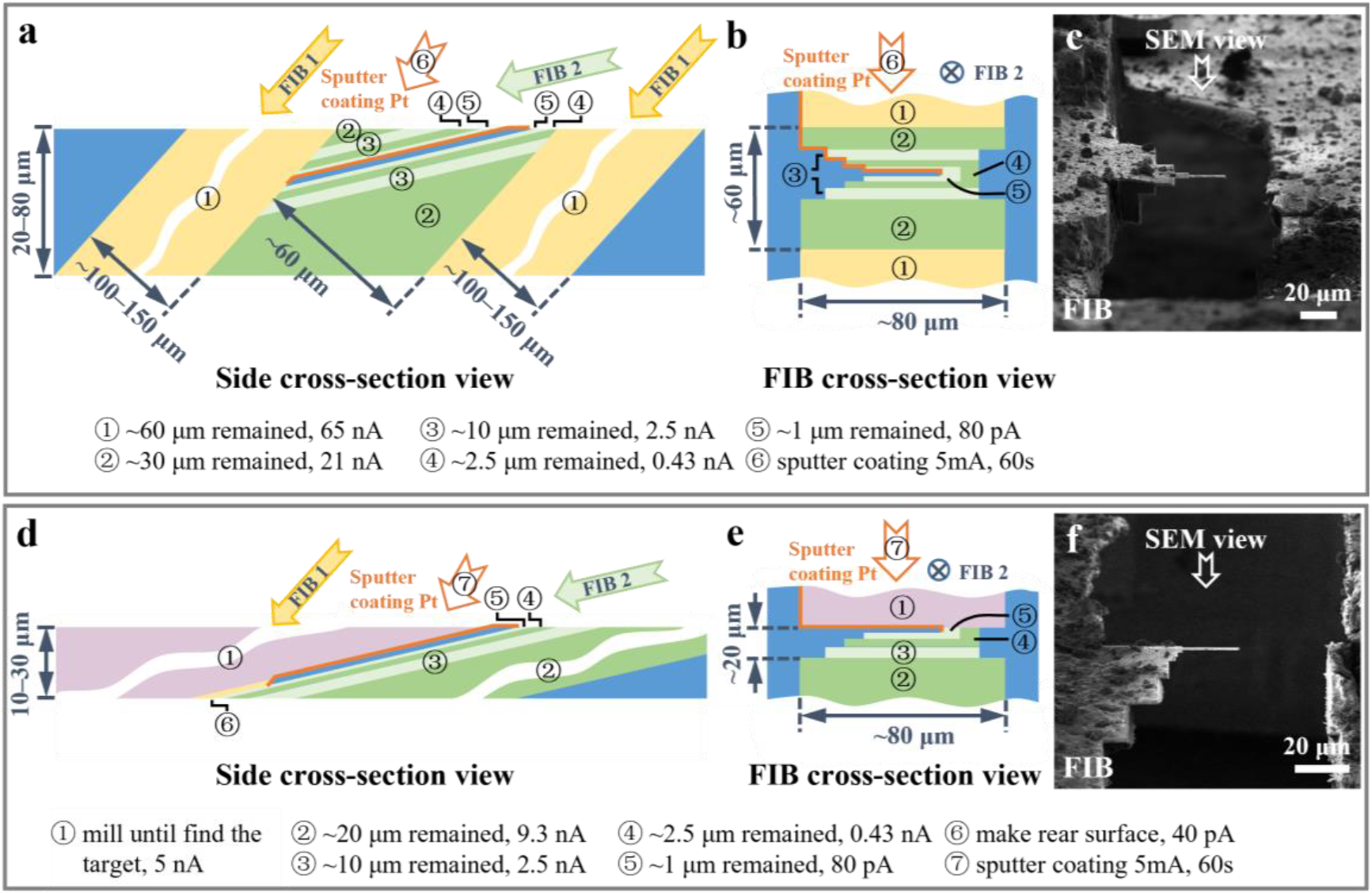
Coarse milling procedure. **a** and **b**, Schematic diagrams of a typical coarse milling procedure viewed from the side (**a**) and FIB cross-section view (**b**). The yellow and green (light and dark green) strips represent the first and the second sub-steps, respectively, with milling along different ion beam directions (arrows labeled with FIB 1 and FIB 2). **c**, Image of a liver sample under FIB view after the coarse milling presented in (**a** and **b**). **d** and **e**, Schematic diagrams of a typical coarse milling procedure skipping the first sub-step to avoid damage on the lamella surface for initial CSEI-based locating. The purple strips represent the volumes removed for CSEI-based locating along the ion beam direction of FIB 2. **f**, Image of a liver sample under FIB view after the coarse milling presented in (**d** and **e**). The strips with different colors and numbers in circles indicate different milling steps and milling parameters (listed at the bottom). The numbers also indicate the sequence of milling and sputter Pt coating (orange edges). The blue volumes and strips are the remaining materials and the final lamella.

CSEI can be used throughout the milling process to determine the milling position. In some cases, CSEI was used ahead of the coarse milling step to find targets sparsely distributed in a large bulky sample. We found that the strong ion beam used in the first milling sub-step could influence a much larger range than the specified milling window and damage the lamella surface generated during CSEI-based locating. Thus, if the target object is close to the surface, the first sub-step should be skipped to avoid damage, and the second sub-step should be applied on another side of the sample (**Fig. 3d-f**). In such a case, we suggest preparing thin samples, typically not thicker than 30 μm, in the pre-sectioning step to avoid long milling times.

The final lamella generated by the coarse milling was typically 1 μm thick. The sputter coating was the last step of the coarse milling process, which deposited a conductive Pt layer on the lamella. This conductive layer was useful for eliminating surface charging during future data collection (discussed later). The coating should cover the front, top, and rear surfaces of the lamella (orange layers in **Fig. 3a** and **Supplementary Fig. 6a**). In the case that the rear surface of the lamella cannot be effectively coated (**Fig. 3d** and **Supplementary Fig. 6b**), the rear surface of the lamella should be milled out using low beam currents (24–40 pA) at a 48° angle to create a sloped surface that can be effectively coated by sputter coating (**Fig. 3d**).

### Fine milling to generate large, thin lamellae

In the fine milling step, we used a beam current as low as 40 pA to finalize the milling process. The lamellae obtained by coarse milling were usually large, typically 20 μm in width and 40–110 μm in length. The primary obstacle to milling such a large lamella to 100–200 nm thickness is the issue of bending (**Supplementary Fig. 7**), which causes milling to fail before reaching the target thickness. To solve this problem, we designed a furrow-ridge structure to enhance the lamella (**Fig. 4**), as well as a corresponding milling workflow (see **Methods**).

**Figure 4.**
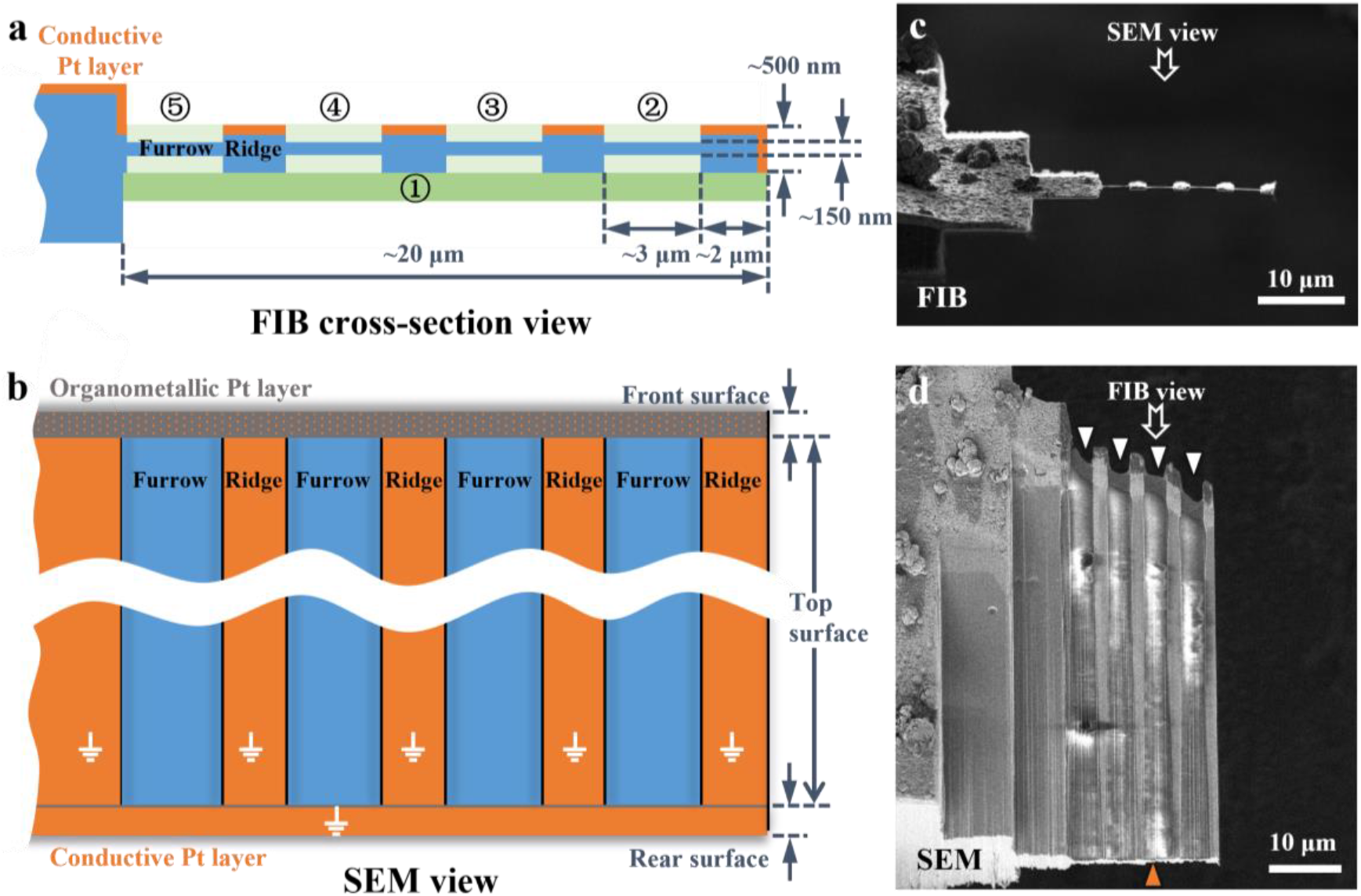
Fine milling procedure and furrow-ridge structure. **a** and **b**, Schematic diagrams of the fine milling procedure under FIB cross-section view (**a**) and SEM view (**b**). The green strips (dark green and light green) represent the volumes removed in the sequence indicated by the numbers in circles. The orange edges and surfaces indicate the conductive Pt layer. The gray surface is the organometallic Pt layer coated by a gas injection system (GIS). The orange dots on the front surface indicate the conductive Pt layer which might be damaged during fine milling. The front, top, and rear surfaces are defined in **Supplementary Fig. 6**. The blue volumes and surfaces are the remaining materials. **c**, and **d**, Finished lamella of a liver sample created by fine milling under FIB view (**c**) and SEM view (**d**), as presented in **a** and **b**, respectively.

The furrow is the thin area to be used for future cryoET data collection, and the ridge is the thick ribbon providing mechanical support and charge reduction using the remaining Pt layer coating applied at the end of the coarse milling step (discussed later). We usually set the furrow to ∼3 µm in width to match the electron beam size during cryoET data collection, and the ridge to ∼2 µm in width. Accordingly, a lamella with a width of 20 µm can accommodate 4 furrow-ridge pairs (**Fig. 4**). This furrow-ridge structure can significantly improve the mechanical strength of the large lamella, and, consequently, the milling success rate.

Excessively heavy ridges can also cause the lamella to bend. Therefore, before milling the furrow-ridge structure, we usually thinned the thick lamella to ∼500 nm from the bottom surface to reduce the weight of the future ridges while maintaining the Pt layer on the top surface (**Fig. 4a**). Then, the furrows were milled successively from the disconnected end to the fixed end of the lamella. In some situations, previously milled furrows bend when milling successive furrows; milling in the correct order improved the likelihood that each furrow was milled to the desired thickness before bending. Fortunately, bending did not obviously influence cryoET data collection and tomogram reconstruction (**Supplementary Fig. 4e-h**).

### Using Pt coated ridges to reduce beam-induced charging

cryoFIB milling of a thick tissue sample exposes a large, hydrated surface that is electrically poor- or non-conductive; therefore, surface charging and the corresponding beam-induced motion are often severe (**Supplementary Fig. 8** and **Supplementary Movie 1**) and inevitably result in the failure of cryoET data collection and tomogram reconstruction. To solve this problem, we performed additional sputter Pt coating applied at the end of the coarse milling step, which covered the ridges with an electrically conductive Pt layer. The Pt layer reduced the non-conductive surface as much as possible and provided a well-grounded metal layer (on the ridges) tightly surrounding the area for cryoET data collection (the furrows). The electrically conductive region has been shown to be essential for neutralizing the surface charge by making the electron beam touch the conductive area during data collection^20, 21^.

Note that the Pt layer on the ridges must be well grounded; this is the reason that the rear surface of the lamella must be sputter Pt coated, as mentioned in the coarse milling step (**Fig. 3** and **Supplementary Fig. 6**). While both the rear and front surfaces of the lamella are Pt coated, the front Pt layer is often damaged by the ion beam, and thus not guaranteed to act as a grounding wire, rendering the rear Pt layer very important (**Fig. 4b**). To demonstrate the role of the conductive Pt layer on the ridges, we performed three comparisons. First, we prepared a lamella with a furrow-ridge structure but without the conductive Pt layer on the ridges. During cryoET data collection, the furrow close to the broken end showed severe motion (**Supplementary Fig. 9a-e**), but the motion was slightly weaker than that of the lamella without a furrow-ridge structure (**Supplementary Fig. 8**). This comparison demonstrates that the mechanical support from the ridge is useful, but not the key factor in reducing motion. Second, we tested a lamella with a furrow-ridge structure on which the conductive Pt layer on the ridge was disconnected from the ground by milling out the rear Pt layer. This lamella exhibited a similar motion as the previous lamella without the Pt layer on the ridges (**Supplementary Fig. 9f-j**), which confirmed that charging was still the problem. Third, we prepared a lamella with a normal furrow-ridge structure, i.e. the ridge had a well-grounded Pt layer. All furrows on the sample showed similar and weak motions. Particularly, the motion of the furrow close to the disconnected end was dramatically reduced by a factor of several tens (**Supplementary Fig. 9k-o**), demonstrating that charging was the key factor causing motion. In the tests above, all of the motion curves were U-shaped, indicating that the motion behaved like a swing around the fixed end of the lamellae.

### Example of milling mouse liver tissue

We used mouse liver tissue to demonstrate the complete milling workflow, with the aim of observing collagen fibrils in the liver. Liver fibrosis results in the excessive accumulation of extracellular matrix (ECM). The structure and degree of crosslinking of collagen fibrils, the major ECM structural protein, have important impacts on liver fibrosis recovery^22, 23^. Therefore, we expected to observe the ultrastructure of collagen fibrils in mouse liver tissue at the early stage of liver fibrosis. The collagen content has been reported to be low in the early fibrotic liver ECM of the carbon tetrachloride (CCl_4_)-induced mouse model^24^, providing an opportunity to test CSEI in locating a sparse target.

Following the protocol described above (see **Method** and **Supplementary Protocol**), fresh liver tissue was fixed in glutaraldehyde for 30 min, sliced using a vibratome (Leica VT1200S, Leica Microsystems Company) set to 20 μm, and frozen using HPF. We randomly milled seven regions on the prepared frozen sample (**Fig. 5a**) and searched for the collagen fibrils using CSEI. Several clusters of collagen fibrils with the characteristic bamboo-like organization were observed in one region (**Fig. 5b**).

**Figure 5.**
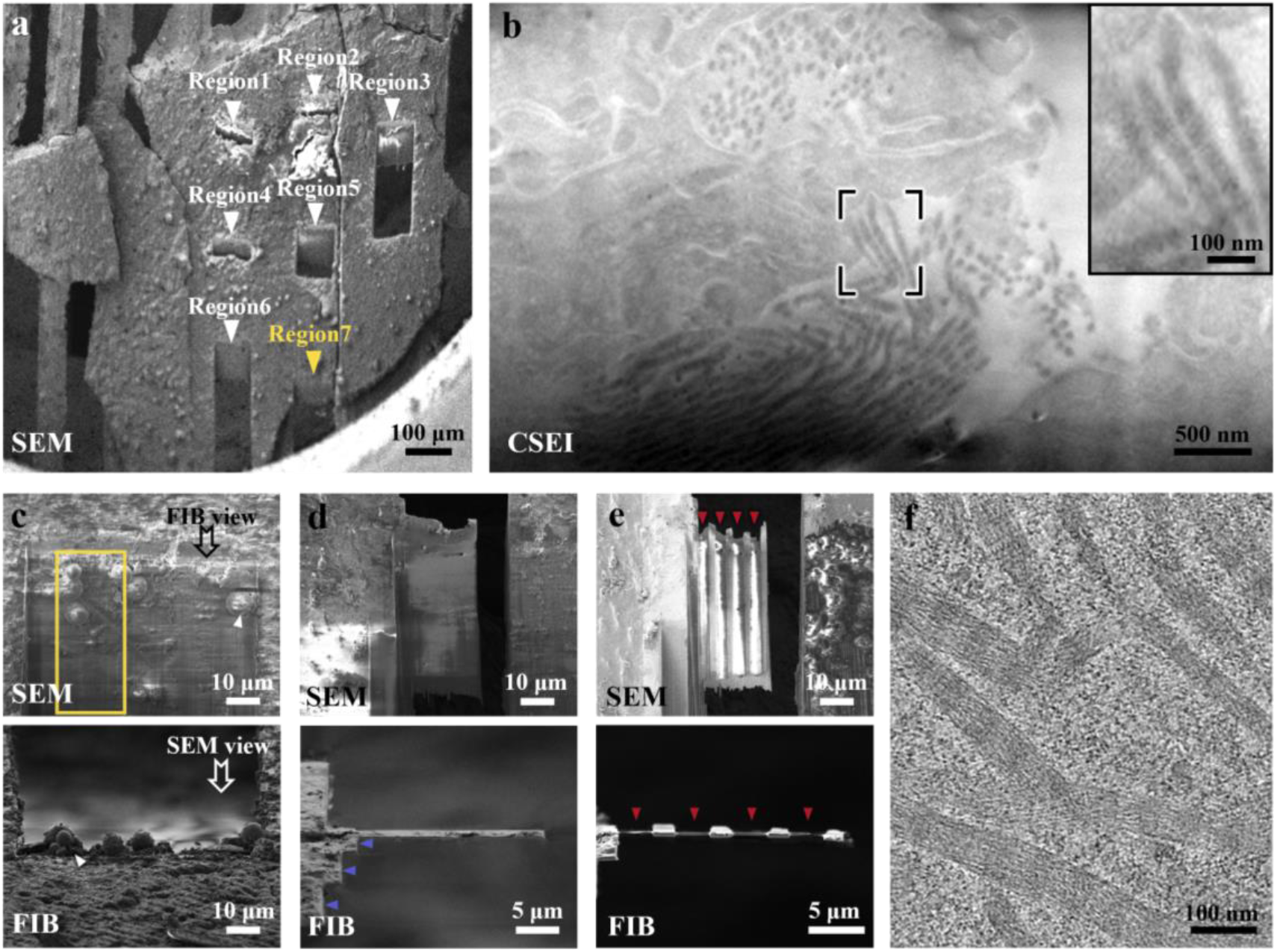
Complete workflow illustrated using a mouse liver sample. **a**, Pre-sectioned and frozen liver slice under SEM view. Seven regions (arrows) were created for CSEI-based locating. **b**, Region 7 in (**a**) was viewed by CSEI, showing the target collagen fibrils. The inset image shows magnified fibrils in the black box. **c**, The top surface of the lamella was milled during CSEI-based locating. The yellow box is the target area. Ice contamination (white arrows) was produced when transferred to another cryoFIB instrument for further milling. **d**, The liver tissue was coarsely milled to reduce the target area to ∼1 µm thick. Blue arrows show the stepped edge on one side of the lamella. **e**, The final lamella after fine milling. Red arrows show the furrows. **f**, A section-view of a tomogram showing collagen fibrils (**Supplementary Movie 2**). The imaging method, SEM or FIB, is labeled in the bottom-left of each panel.

Subsequently, around the collagen fibrils, we generated large lamellae by coarse milling (**Fig. 5c-d** and **Supplementary Fig. 4a-d**), and further milled them into thin lamellae with four furrow-ridge pairs using the fine milling protocol (**Fig. 5e** and **Supplementary Fig. 4e-f**). Using a 300 kV cryo-electron microscope, we clearly observed collagen fibrils widely distributed in the lamella (**Supplementary Fig. 4g**). We then collected tilt series data and performed tomography reconstruction. The tomograms showed high contrast features of the D-period of collagen fibrils^22, 25^, and the lamella thickness was approximately 130 nm (**Fig. 5f, Supplementary Fig. 4h**, and **Supplementary Movie 2**). In another sample, we obtained a thinner lamella of ∼100 nm thick, in which more cellular ultrastructures were observed, including mitochondria and endoplasmic reticulum (**Supplementary Fig. 10** and **Supplementary Movie 3**).

## Discussion

In this study, we proposed a complete workflow for milling large tissue samples using cryoFIB, which involved pre-sectioning, on-the-fly locating, and rapid milling. This workflow required only the basic functions of a cryoFIB instrument, including secondary electron imaging and focused ion beam milling. CSEI provides a convenient way to accurately locate the samples and eliminate reliance on fluorescence labels. More importantly, CSEI is very reliable on various surfaces created by cryoFIB milling, which allows for constant imaging and the determination of the 3D position of target objects during milling. Together with our high-efficiency cryoFIB milling strategy, preparing the thin lamella starting from a large tissue has become achievable. Finally, we demonstrated the entire workflow using a mouse liver and obtained high-quality tomograms of the collagen fibers sparsely distributed in the liver sample.

The bottleneck for thinning tissue is efficiency. We developed two strategies to solve this problem. The first strategy involved coarse milling using an ion beam current as high as 65 nA to rapidly remove the volume surrounding the target object. The second strategy was to generate a large lamella with a width and length of tens of micrometers, allowing us to expose a large sample area for cryoET analysis. Combining the two strategies, the milling efficiency we achieved when starting from a large tissue sample was on par with that achieved when starting from a thin single-cell sample. The major problem currently affecting the success rate is ice contamination, which occurs during the transfer process after milling and can significantly reduce the area suitable for cryoET data collection. Efforts to reduce ice contamination are still necessary.

The furrow-ridge structure was the key to obtaining high-resolution cryoET data. This structure allowed us to successfully obtain quality lamellae as thin as 100 nm. Lamella thinness is an essential factor for achieving sub-tomogram averaging at near-atomic resolution. Furthermore, the well-grounded conductive Pt layer on the ridge provided an effective way to relieve the beam-induced motion problem disturbing cryoET data collection and subsequent analysis. These innovations improved the success rate and accuracy of tracking and tomography alignment of series-tilting images, as well as the quality of frames collected at high tilt angles.

In summary, our workflow for tissue sample preparation eliminates the major obstacles in the preparation of large tissue samples for cryoET and efficiently broadens the applicable range of cryoET to nearly any large biological sample.

## Supporting information

Supplementary Protocol

## Contributions

X.L. initialized the project. X.L. and S.W. designed the fixation, pre-section, and HPF experiment. H.Z. designed the cryoFIB milling strategy. X.L., S.W., and H.Z. designed the CSEI experiment. S.W and H.Z performed all the experiments. Y.J. assisted with the cryoFIB milling and imaging on the CrossBeam 500 instrument. W.C., X.Y., and H.Y. established a liver fibrosis mouse model. X.L, S.W, and H.Z wrote the manuscript. All authors revised the manuscript.

## Acknowledgments

This work was supported by funds from Tsinghua-Peking Joint Center for Life Sciences, Beijing Frontier Research Center for Biological Structure, and Advanced Innovation Center for Structural Biology. We thank Ying Li for technical assistance in the HPF experiments, Weilin Huang for assisting the image processing, Bingyu Liu for assistance in operating Amira 20.2, Xiuduan Xu for providing liver tissue of non-fibrosis mouse, Hao Wang and Xiaomin Li for technical assistance in cryoET data collection. We acknowledge ZEISS Microscopy Customer Center, Beijing lab for providing SEM-FIB facilities. We acknowledge the Tsinghua University Branch of China National Center for Protein Sciences Beijing for providing facility support.

## Data availability

The tomograms of collagen fibrils have been deposited into the Electron Microscope Data Bank (EMDB) with the accession code EMD-33910 for Figure. 5f and the accession code EMD-33911 for Supplementary Figure. 4h. The tomograms of mice liver tissue shown in Supplementary Figure. 10a has been deposited into EMDB with the accession code EMD-33912, and the corresponding one processed by IsoNet was deposited with the accession code of EMD-33913.

