## Supplementary Materials for "CryoFIB milling large tissue samples for cryo-electron tomography"

### Methods

#### $CCl_4$-intoxicated liver fibrosis mouse model

Adult C57BL/6 mice (8 weeks, male) were purchased from Cyagen Biosciences (Suzhou, China) and acclimatized in cages for 3 days. The liver fibrosis mouse model was established by intraperitoneally injecting 12.5% of $CCl_4$ (Innochem, China) in mineral oil (1/7, v/v) at a dose of 0.01 ml/g body weight twice a week for up to 8 weeks (for early fibrosis). Healthy mice only received isovolumic mineral oil in the same manner and were used as controls. All mice were housed and bred at 23±2℃, under a 12-hour light-dark cycle with standard chow and water *ad libitum*. Mouse studies were approved by the Ethics Committee of Beijing Friendship Hospital, Capital Medical University, and carried out following the stated guidelines.

#### Liver tissue sample acquisition and vibratome sectioning

Mice were euthanized using excess pentobarbital sodium. Liver tissues were dissected and cut into small cuboids (with an edge length of 1–3 mm) with a razor blade. Subsequently, the liver tissue cuboids were washed 2–3 times in cold phosphate buttered saline (PBS) and submerged in 2.5% glutaraldehyde (Electron Microscopy Sciences) for 30 min at room temperature. Subsequently, liver tissue cuboids were washed 2–3 times in cold PBS and submerged in 4% liquid agarose (AGARSE II, 0815-5G, Amresco) at approximately 40℃. After agarose solidification, the agarose was mounted on the sample stage of a vibratome (Leica VT1200S, Leica Microsystems). The buffer tray of the vibratome was filled with cold PBS. The vibratome was set to section at a thickness of 10–50 µm (sectioning frequency: 85 Hz; amplitude: 1 mm; sectioning speed: 0.5 mm/s). The liver slices were then carefully transferred to cold PBS using a brush.

#### High-pressure freezing of liver tissue

The liver slices (immersed in PBS) were picked up using a Perfect Loop (Diatome, DZ8) and loaded onto glow-discharged grids coated with lacey carbon film (parallel bars, Cu, 150 mesh, Zhongjingkeyi Technology, AG150P) (**Fig. 1b**). Next, grids with tissue slices were blotted on filter paper to remove excess liquid (~5 s) and transferred into aluminum carriers (6 mm aluminum specimen carrier Type A, 200 µm recess, Leica Microsystems), which were previously filled with 2-methylpentane (Sigma, M65807). The aluminum carriers were then covered with sapphire discs and immediately transferred to a high-pressure freezer (Leica HPM100, Leica Microsystems). After cryo-freezing, the aluminum carriers were transferred into a cryo-ultramicrotome chamber (Leica EM UC7+FC7, Leica Microsystems) at -150℃ to melt the frozen 2-methylpentane (melting point, -154℃). Thereafter, grids were separated from the aluminum carriers and placed in the cryo-ultramicrotome chamber for ~20 min to remove excess 2-methylpentane. Finally, grids were transferred to grid boxes and stored in liquid nitrogen.

#### Preparation for cryoFIB milling

The frozen grids containing tissue slices were mounted into Autogrids (Thermo Fisher

Scientific). The Autogrids were then mounted onto a custom-made sample shuttle. Note that the direction of the Autogrid should be aligned such that the grid bars point toward the incident direction of the ion beam (see **Supplementary Protocol**).

The shuttle was transferred to the prep-stage in the prep chamber of the cryo-transfer system (PP3010T, Quorum Technologies), which was pre-cooled to -180℃. The sample was sublimated under high vacuum in the prep chamber at -150℃ for 30–90 min to ensure complete removal of 2-methylpentane and exposure of the tissue bulk. After sublimation, the temperature of the prep-stage was set back to -180℃. The shuttle was then transferred to the cryo-stage in the SEM chamber (Helios Nanolab G3 UC, Thermo Fisher Scientific) and elevated to a position such that the grid was 8 mm from the SEM tip. The acceleration voltage and beam current of the SEM were set to 2 kV and 0.4 nA, respectively. The organometallic Pt gas in the gas injection system (GIS) was preheated to 42℃. Then, electron-beam-induced organometallic Pt deposition was performed for 40 sec with a continuous electron beam scanning over the entire coated area at a magnification of 100×. To improve the conductivity, the shuttle was transferred back to the prep chamber for sputter coating (5 mA, 60 s) to cover the organometallic Pt layer with a thin conductive Pt layer.

**CSEI for locating**

The Autogrids were mounted onto a Zeiss-customized shuttle and transferred to a Zeiss Crossbeam 550 FIB-SEM (Carl Zeiss Microscopy, Oberkochen, Germany) using a QUORUM PP3010Z transfer system. Throughout the imaging, the samples were kept below -170℃. The cryo-stage was tilted to 8° to produce lamellae at a shallow angle (18° relative to the grid plane). A section surface of 80 μm in width was milled by cryoFIB with the following settings: 30 kV acceleration voltage, 5–7 nA ion beam currents. After milling, each milled surface was simultaneously imaged by CSEI using the in-lens and in-chamber detectors at 3 kV acceleration voltage. The SEM imaging settings included an electron beam current of 50 pA, image size of 1024×768 pixels, dwell time of 1.8 μs (scan speed of 5), and repetitive scans of 20 times. The in-lens/in-chamber mixed detection was performed using the smartSEM (Carl Zeiss Microscopy) with a mixing ratio between 0.5–0.7.

**CryoFIB milling procedure**

A dual-beam FIB/SEM (Helios Nanolab G3 UC, Thermo Fisher Scientific) with a cryo transfer system (PP3010T, Quorum Technologies) was used for lamella preparation. The angle between the electron beam (for SEM) and ion beam (for cryoFIB) was 52°. A 2 kV accelerating voltage and a 50 pA electron beam current were used for SEM imaging, and a 30 kV accelerating voltage and a 40 pA ion beam current were used for cryoFIB imaging. Throughout the cryoFIB milling process, the samples were kept below -170℃ and the system vacuum pressure was approximately $1.5\times10^{-4}$ mbar.

The entire cryoFIB milling procedure consisted of two steps: coarse milling and fine milling (see **Supplementary Tables 1 and 2** for typical parameters). The coarse milling procedure was performed in two sub-steps (see **Supplementary protocol** for additional guidance). In the first sub-step, the sample stage was tilted such that the incident direction of the ion beam and the grid plane formed a high angle of 48°. CryoFIB milling using an ion beam current of 65 nA was performed to remove sample volume in two rectangular windows (80×100μm under FIB view) using the Rectangle Pattern (named in the FIB user interface) at the front and back ends of the target position, resulting in a slab of 60 μm in thickness under FIB view (**Supplementary Figs. 2 and 3a-b**). In the second sub-step, the FIB incident angle was adjusted to the final milling angle of the lamella (between 13–18°) (**Supplementary Fig. 3d**). The second sub-step of the coarse milling process was performed by stepwise reduction of the ion beam current (from 21 nA, 9.3 nA, 2.5 nA, 790 pA, to 80 pA) and the width of rectangle window (from 70 μm, 50 μm, 30 μm, 25 μm, to 20 μm) to create a stepped edge around the lamella. The lamella was thinned to approximately 1 μm in thickness by the stepwise movement of the rectangular windows closer to the final lamella (**Supplementary Fig. 3e-g). For the lamellae previously located by CSEI, only the second sub-step of the coarse milling process was performed on the bottom surface using stepwise reducing ion beam current (from 9.3 nA, 2.5 nA, 790 pA to 80 pA) (**Supplementary Fig. 4**). The width of the rectangular milling window was gradually reduced to create a stepped edge on one side of the lamella. The rear of the lamella was milled out using a 40 pA ion beam current at a 48° angle. In addition, one side of the lamellae was disconnected from the bulky sample during coarse milling. Lamellae were transferred to the prep chamber for sputter coating (5 mA, 60 s). Fine milling was performed using a 40 pA ion beam current to prepare the final lamellae with 4 furrow-ridge pairs. The furrows and ridges had widths of 3 μm and 2 μm, respectively (**Supplementary Fig. 3i).**

**CryoEM imaging and cryoET data collection**

After cryoFIB, the Autogrids were loaded into a Krios Cassette, ensuring that the milling direction of lamellae was perpendicular to the tilt axis for cryoET data collection. The lamellae were imaged using an FEI Titan Krios microscope (Thermo Fisher Scientific) operated at a voltage of 300 kV and equipped with a Quantum post-column energy filter (Gatan) and a K3 Summit direct electron detector (Gatan). Image acquisition was controlled using SerialEM[26]. Montages of the whole lamella were acquired at 2250× magnification. Two magnifications were used for cryoET data collection. The collagen fibril tomograms (**Fig. 5f** and **Supplementary Fig. 5h**) were acquired at a magnification of 33,000× (pixel size of 2.12 Å). The tomogram shown in **Supplementary Fig. 10a** was acquired at a magnification of 19,500× (pixel size of 3.65 Å). Tilt series were acquired using a bidirectional tilt scheme starting from -20° relative to the lamella plane (i.e. from -7° to 67°, then from -9° to -47° for a 13°-tilted lamella, and from -2° to 68°, then from -4° to -46° for an 18°-tilted lamella) with a tilt angular step of 2°. The target defocus was in the range of -4 to -6 μm. Micrographs with 8 frames were recorded with a 0.075 s per frame exposure time at each tilt angle, resulting in an electron dose of 2 $e^-/Å^2$. The total accumulated electron dose for a tilt series is ~116 $e^-/Å^2$. During cryoET data collection, the electron beams for recording/tracking/focusing illumination were controlled to ensure the touch with the ridges to eliminate charging. The tracking/focusing area was selected in an adjacent furrow close to the recording area.

**Image processing**

Dose fractionated images recorded by the Gatan K3 Summit camera were motion-corrected using MotionCorr2[27]. All tilt series were aligned using patch tracking and reconstructed by Simultaneous Iterative Reconstruction Technique (SIRT) using the IMOD software package (version 4.9.12)[28]. The tomogram was enhanced using IsoNet[29]. Tomogram segmentation and 3D visualization were performed using Amira 20.2 (Thermo Fisher Scientific, Mercury Computer Systems).

**Motion analysis**

Different types of lamellae were prepared using a dual-beam FIB/SEM (Helios Nanolab G3 UC, Thermo Fisher Scientific). The first type was a lamella without the furrow-ridge structure. The lamella was milled to ~150 nm thickness. The second type was a lamella with a furrow-ridge structure but without the conductive Pt layer on the ridges. The lamella contained 4 furrow-ridge pairs, but no additional sputter Pt coating was applied following coarse milling. The third type was a lamella with a furrow-ridge structure, but the conductive Pt layer on the ridge was disconnected to the ground by milling out the rear Pt layer at a 48 ° milling angle after sputter coating.

The above lamellae and a lamella with a normal furrow-ridge structure were transferred to the FEI Titan Krios electron microscope (ThermoFisher Scientific). Tilt series were collected at a magnification of 33,000× with a pixel size of 2.12 Å and using a bidirectional tilt scheme starting from -20 ° relative to the lamella plane (i.e. from -7 ° to 67 °, then from -9 ° to -47 ° for a 13 °-tilted lamella, and from -2 ° to 68 °, then from -4 ° to -46 ° for an 18 °-tilted lamella) with a tilt angular step of 2 °. The target defocus was set in a range of -4 to -6 μm. Micrographs with 8 frames were recorded with a 0.075 s per frame exposure time at each tilt angle, resulting in an electron dose of 2 $e^-/Å^2$. Some micrographs at high tilt angles were unable to be collected due to the failure of tracking caused by severe motion, resulting in a varying number of micrographs in a tilt series. MotionCor2[27] was used for whole-frame motion correction. Drift at each angle was measured by calculating and summing the displacement between adjacent frames of the 8-frames micrograph.

### Supplementary Figures and Legends

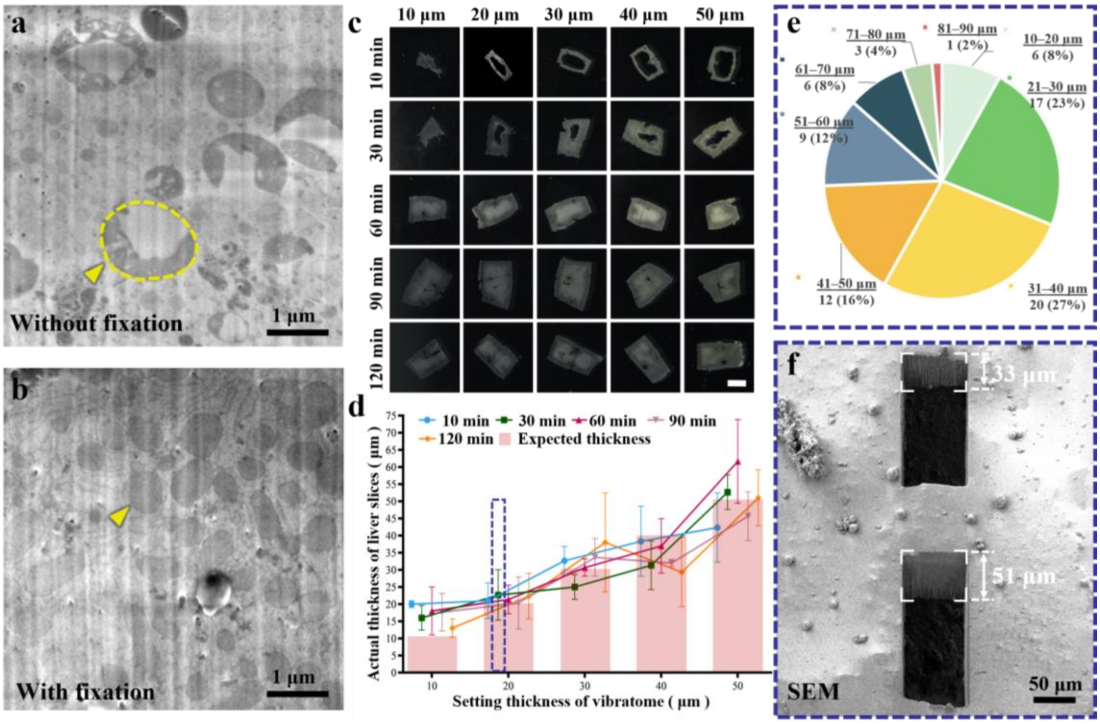


**Supplementary Figure 1. Chemical fixation and pre-sectioning of liver tissue. a** and **b,** CSEI of a frozen hydrated liver tissue sample processed without (**a**) or with (**b**) fixation ahead of pre-sectioning. The pre-sectioning process required approximately 15 min and caused some cell damage. The cellular contents released from broken cells may have influenced adjacent liver cells in the unfixed sample, resulting in mitochondria with a tumid feature (yellow arrow and dashed circle in **a**). 2.5% glutaraldehyde fixation for 30 min reduced this effect, as indicated by the compact mitochondria shape (yellow arrow in **b**). **c,** Photos of liver tissue slices prepared with different pre-sectioning thicknesses (labeled at the top) and fixation times (labeled on the left) using 2.5% glutaraldehyde. The liver tissues fixed for 10 min exhibited a large broken central region. The size of the broken region was reduced with increasing fixation time. Liver slices fixed for 60–120 min were intact. The brightness of the central region also gradually darkened with increasing fixation time, which may indicate more glutaraldehyde diffusion into the central region. **d.** Statistical analysis of the actual thickness of liver slices prepared using a vibratome. The expected thickness is the setting thickness of the vibratome, shown as pink bars. Each data point was measured from at least three samples and shown as the mean and standard deviation (error bars). The large error bars indicate a large variation between the setting thickness in vibratome and the actual slice thickness. Data points measured on samples with different fixation times are shown with different colors. The samples from one of the conditions (dark-blue dotted box) are individually analyzed in (**e** and **f**). **e,** Statistical distribution of the sample thickness measured from the 74 samples included in the condition labeled by the blue dotted box in (**d**). **f,** Two windows penetrating the sample were milled by FIB along an incident angle of 48$^{o}$ relative to the sample surface. Under SEM view, the walls of the two windows exhibited differing heights of 33 μm and 51 μm, respectively, demonstrating the uneven thickness of the liver slices prepared by the vibratome.

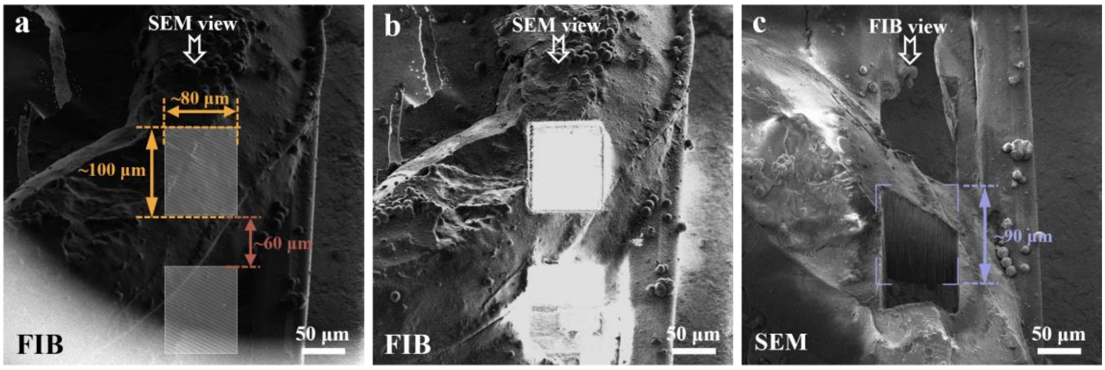


**Supplementary Figure 2. Example of the first sub-step of coarse milling. a**, Two windows of ~80×100 $\mu m^2$ separated by 60 µm were planned for milling under FIB view. The ion beam was along a large incident angle of 48 °relative to the grid plane. **b**, The two windows were milled using a 65 nA ion beam current, imaged under FIB view. **c**, The SEM view of the two milled windows. The walls of the window were ~90 µm in height under SEM view. Considering the angle between the electron beam (for SEM) and the ion beam (for cryoFIB) was 52 °, the actual thickness of the sample was ~84 µm.

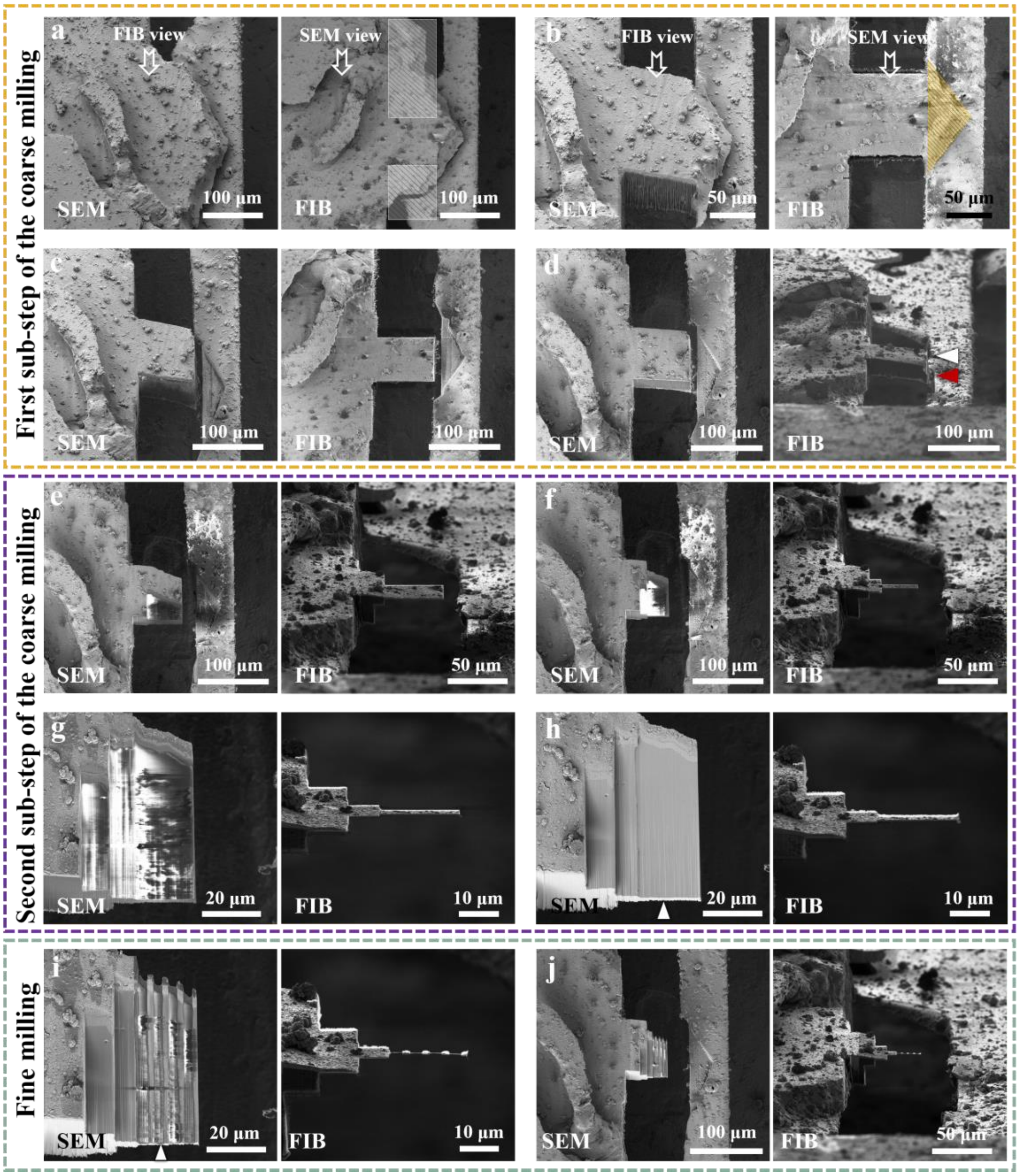


**Supplementary Figure 3. Example of the complete milling procedure.** A frozen hydrated mouse liver sample was used for this example. **a,** The sample was tilted such that the ion beam milled along an incident angle of 48 °. Two windows were planned under the FIB view for the first sub-step of coarse milling. A pair of images under SEM view (left) and FIB view (right) are shown for all panels. **b**, The two windows were milled using a 65 nA ion beam current at an incident ion beam angle of 48 °, as specified in (**a**). **c,** An edge of the sample (yellow triangle area in **b**) was removed using a 65 nA ion beam current at an incident ion beam angle of 48 °. The edge was loosely attached to the grid bar and may have been unstable once the target area was thinned; therefore, removing the edge ahead of further milling may have helped to maintain the stability of the target area. **d,** The sample viewed after adjusting the incident ion beam angle to 13 ° relative to the grid plane. This angle was maintained in subsequent milling. The target milling position should be chosen on the original sample surface covered by an organometallic Pt layer (white arrow) but not the freshly milled surface (red arrow). **e,** The sample was milled to 10 μm thick in the second sub-step of coarse milling using a beam current of 2.5 nA. **f**, The sample was milled to 2.5 μm thick in the second sub-step of coarse milling using a beam current of 0.43 nA. **g**, The sample was milled to 1 μm thick in the second sub-step of coarse milling using a beam current of 80 pA. This was the final thickness following coarse milling. **h,** The sample was processed by sputter coating using a 5 mA current for 60 s. The rear surface of the lamella is indicated by a white arrow. **i,** The final lamella with the furrow-ridge structure. The lamella had a width of 20 μm, length of ~50 μm, and thickness of ~130 nm, and contained 4 furrow-ridge pairs. The furrows and ridges have widths of 3 μm and 2 μm, respectively. The bright edge on the rear surface of the lamella (white arrow) was the metallic Pt layer generated during sputter coating. **j,** The overall view of the lamella under low magnification.

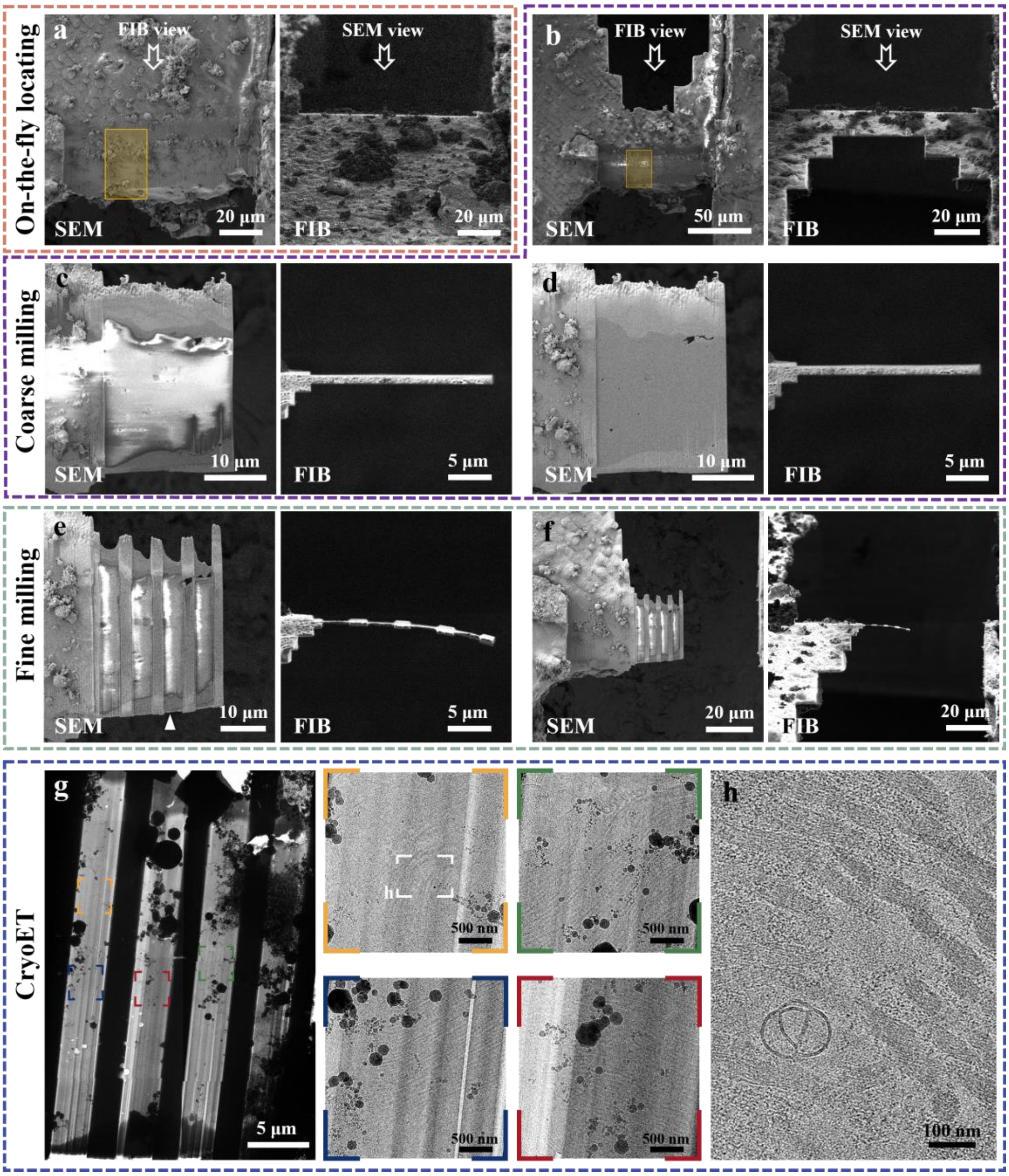


**Supplementary Figure 4. Example of a complete workflow to observe collagen fibrils in mouse liver tissue.** This figure shows the cryoFIB milling procedure on one side of the lamella after the objects of interest, collagen fibrils, were located by CSEI. Note that the first sub-step of coarse milling was skipped in order to avoid radiation damage caused by the strong ion beam. **a,** The surface milled for CSEI locating. The target region containing collagen fibrils is marked by the yellow box. Images under SEM view (left) and FIB view (right) are shown in panels (**a**) through (**f**). **b,** The lamella milled to 10 μm thick during coarse milling. The target region containing collagen fibrils is marked by the yellow box. **c,** The lamella further milled to 1 μm thick, which was the final thickness following coarse milling. Note that the right side of lamella was disconnected from the bulky sample by a milled gap of 30 μm. **d,** The sample was processed by sputter coating. **e,** The final lamella with 130 nm thickness, generated after fine milling. The edge on the rear surface of the lamella (white arrow)

was the metallic Pt layer generated during sputter coating. **f,** The overall view of the lamella under low magnification. **g,** CryoEM image of the lamella under low magnification. The collagen fibrils (colorful boxes and corresponding magnified images shown on the right) in different areas of the lamella were observed. **h,** A slice view of a tomogram collected from the labeled region indicated by the white box in (**g**), which shows characteristic features of the collagen fibrils (**Supplementary Movie 2**).

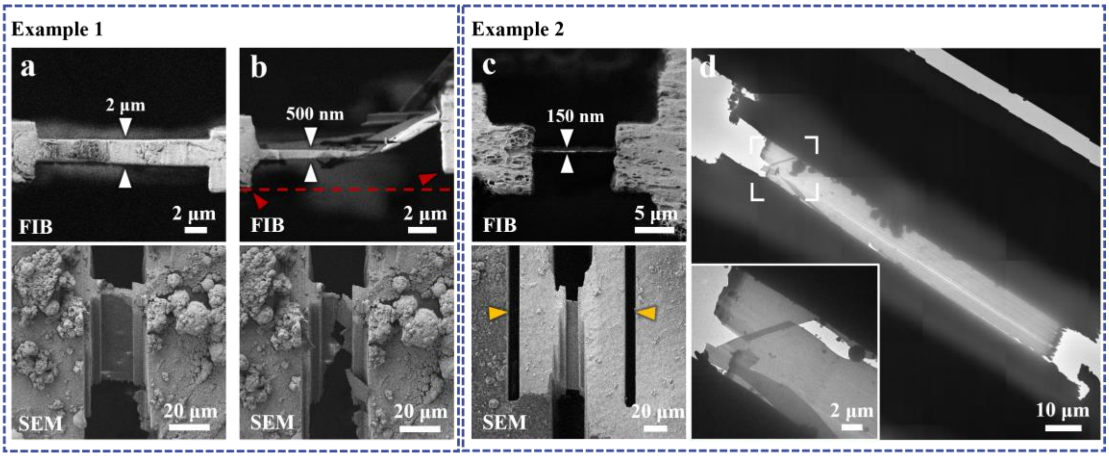


**Supplementary Figure 5. Examples of broken lamellae.** This figure demonstrates the importance of disconnecting one side of the lamella from the thick bulky sample to avoid breakage. **a,** A half-finished lamella of 2 µm thickness. **b,** The lamella shown in (**a**) broke when further milled to 500 nm thickness. The FIB view shows an offset between the left and right edges of the lamella (red arrows). **c,** Another lamella with 150 nm thickness. Two micro-expansion joints (yellow arrows) were created in order to release the inner stress[30]. The images under the FIB view (top) and SEM view (bottom) are shown in panels (**a**) through (**c**). **d**, An image showing that the lamella was broken after being transferred to an electron microscope for cryoET data collection. The inset image shows the magnified area in the white box viewed from a different tilt angle. This indicates that the micro-expansion joint is often not sufficient for samples with large thicknesses.

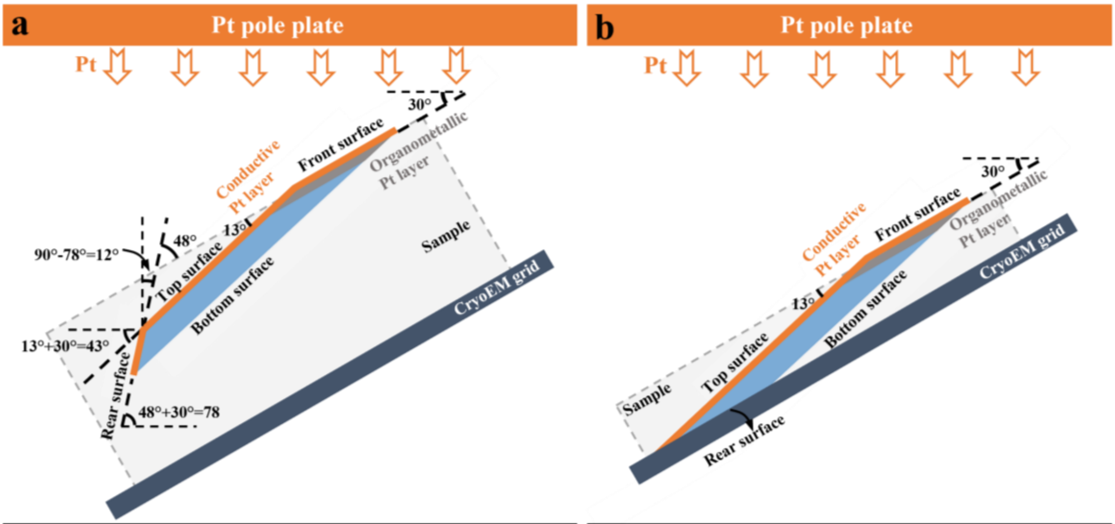


[redacted]

**Supplementary Figure 6. Schematic diagrams of the sputter coating performed after coarse milling. a.** The sputter coating and angular configuration of a sample inside the chamber for sputter coating, corresponding to **Fig. 3a**. The metallic Pt is deposited along the direction from the top Pt pole plate of the sputter device to the bottom. The cryoEM grid is mounted with a tilt angle of 30°. The front surface of the lamella is the top surface of the original sample and is parallel to the cryoEM grid plane. The angle between the top and the front surface of the lamella is determined by FIB direction 2, shown in **Fig. 3a** (13° in this case). The bottom surface of the lamella is parallel to the top surface. The angle between the rear surface and the front surface of the lamella is determined by FIB direction 1, shown in **Fig. 3a** (48° in this case). The angle between the rear surface of the lamella and the horizontal plane is 78° and must not exceed 90°, otherwise, the rear surface will be parallel to the direction of the Pt coating or even shadowed, in which case it cannot be coated. **b,** The sputter coating and angular configuration of a sample after skipping the first sub-step of coarse milling inside the chamber for sputter coating, corresponding to **Fig. 3d**. The rear surface of the lamella is the bottom surface of the original sample, i.e., not facing the Pt source, and therefore, cannot be coated. Therefore, the milling step labeled with the number 6 shown in **Fig. 3a** is required to create a proper rear surface prior to sputter coating. All diagrams were drawn in side view. The metallic Pt element is colored orange, the organometallic Pt layer is colored dark grey, the lamella is colored light blue, the cryoEM grid is colored navy blue and the bulky sample is represented as light gray volume.

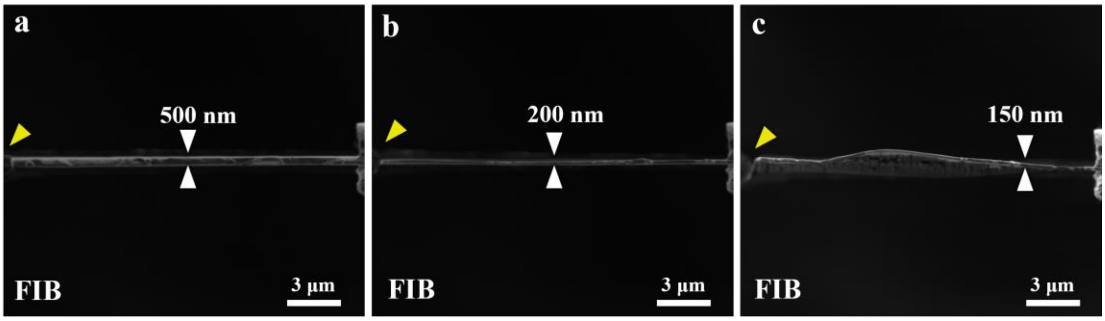


**Supplementary Figure 7. Testing a lamella without the furrow-ridge structure.** The left side of the lamella is disconnected from the bulky sample (yellow arrows). **a,** Lamella of 20 μm in width was thinned to approximately 500 nm. **b,** The lamella was further thinned to about 200 nm, which resulted in slight bending. **c,** The bending became more severe when the lamella was further thinned. The thinnest region of the lamella is approximately 150 nm, and other regions cannot be accurately measured due to bending.

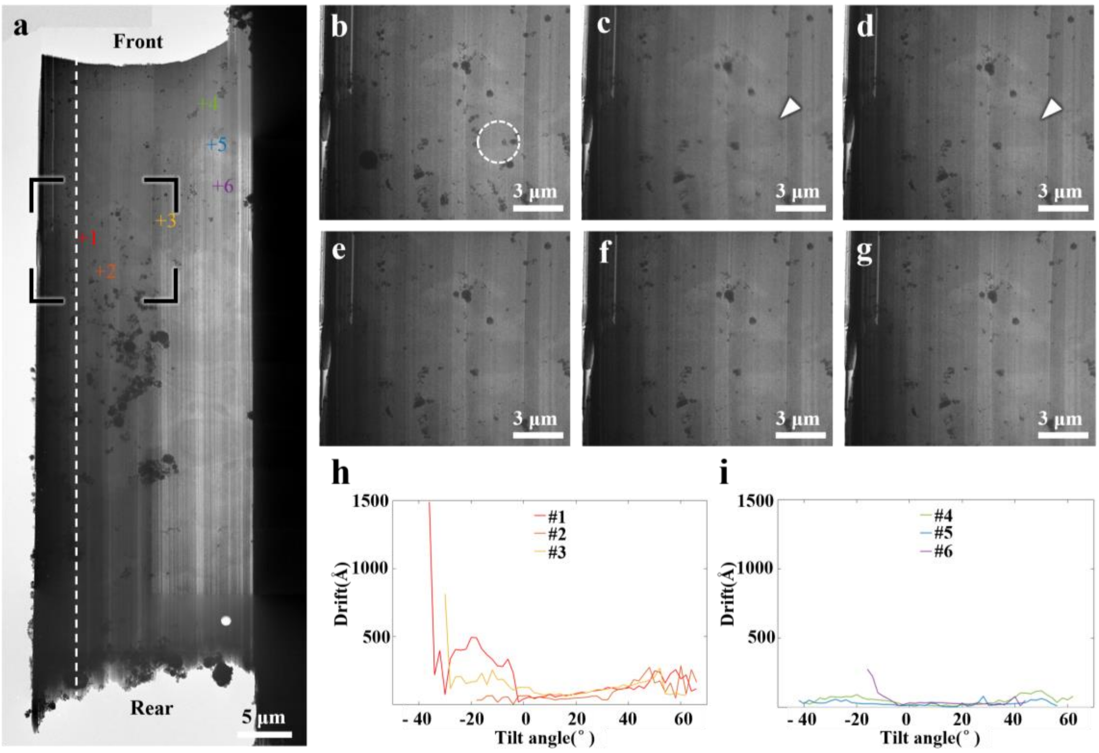


**Supplementary Figure 8. Observation of the charging and beam-induced motion of a lamella without the furrow-ridge structure. a,** A representative cryoEM image of a large lamella (20×60 μm). A region (black frame) was selected for observation. The front and rear surfaces of the lamella are marked. The areas used for tilt-series data collection are shown in colorful crosses labeled with numbers. The motion at the areas very close to the disconnected end of the lamella (on the left of the white dotted line) was so strong that the tracking at all tilt angles failed, hence no cryoET data was collected at these areas. **b,** A cryoEM image taken at the position indicated by the black frame in (**a**) before the exposure. The exposure was performed at a magnification of 33,000× and a dose of 2 $e^{-}/Å^{2}$. The illumination area of exposure is indicated by the white dashed circle. **c-g,** Continuous imaging of the same position as in (**a**) after the exposure (see also **Supplementary Movie 1**). An electron footprint[20] (white arrow) and image distortion (see also **Supplementary Movie 1**) is built up by the charge accumulation in the illuminated area and gradually faded out during continuous imaging at low magnification. **h-i,** Curves of the measured drift (motion) against the tilting angles of cryoET data collection. The motion was measured in the regions labeled with colorful numbers in (**a**). A large motion was observed in regions 1–3 near the disconnected edge. The motion in regions far from the edge, 4–6, was less. Nevertheless, the motion in all six regions was quite severe. In some curves, some data points at high tilt angles were missed or severely fluctuated due to the failure of tracking in data collection.

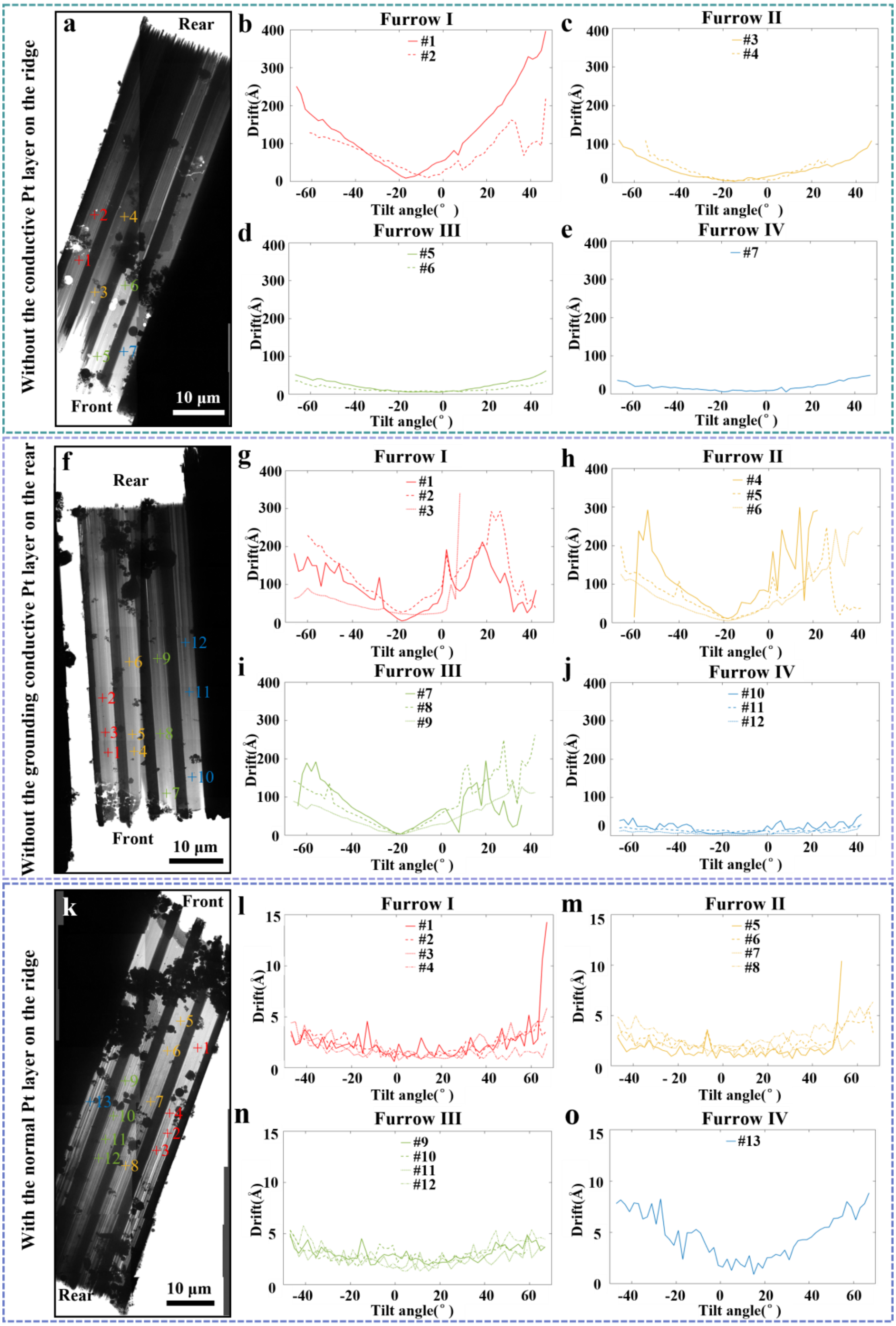


**Supplementary Figure 9. Motion resulting from different Pt layers on the furrow-ridge structure.** Three lamellae were tested and compared**. a,** A low-magnification cryoEM image of a lamella without the conductive Pt layer on the ridge. **b-e,** Curves of the measured drift (motion) against the tilting angles of cryoET data collection. The motion was measured in several regions labeled with colorful numbers in (**a**). The four furrows from the disconnected edge were defined as furrows I–IV in sequence. The colors, red, yellow, blue, and green, were used to indicate positions in furrows I–IV, respectively. For cryoET data collected in different furrows, drift severity was more severe closer to the disconnected edge. **f,** A low-magnification cryoEM image of a lamella with a conductive, but not grounded, Pt layer on the ridge. **g-j,** Curves of the measured drift (motion) against the tilting angles of cryoET data collection. Similar to (**b**) through (**e**), the motion was measured in several regions labeled with colorful numbers in (**f**). The motion behaviors shown in (**g**) through (**j**) are similar to those of (**b**) through (**e**). **k,** A low-magnification cryoEM image of a lamella with a normal Pt layer on the ridge. **l-o,** Curves of the measured drift (motion) against the tilting angles of cryoET data collection. Similar to (**b**) through (**e**), the motion was measured in several regions labeled with colorful numbers in (**k**). The motion measured in different furrows was significantly reduced, and similar in magnitude (below 15 Å). In some curves, some data points at high tilt angles were missed or severely fluctuated due to the failure of tracking in data collection.

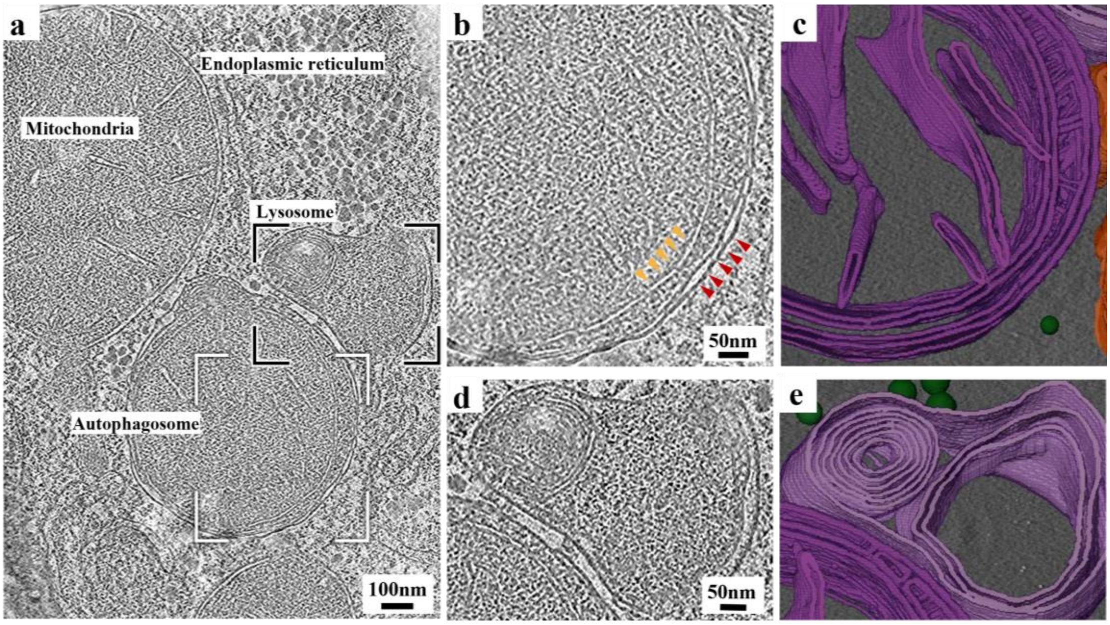


**Supplementary Figure 10. Tomogram of a liver tissue lamella.** A movie of the tomogram is shown in **Supplementary Movie 3**. **a,** Representative section view of a tomogram obtained from a lamella of healthy mouse liver tissue. **b,** Magnified view of an autophagosome labeled by the white box in (**a**). The double-membrane of the autophagosome is indicated by red arrows. The double-membrane of a mitochondrion wrapped inside the autophagosome is indicated by yellow arrows. **c,** 3D visualization of the membrane structure in (**b**). **d,** Magnified view of a lysosome labeled by the black box in (**a**). **e,** 3D visualization of the membrane structure in (**d**).

### Supplementary Tables

**Supplementary Table 1. Typical milling settings and milling times.** All data were based on samples of ~50 µm thickness.

| | Coarse milling | | | | | | | Fine milling | |
|---|---|---|---|---|---|---|---|---|---|
| | **Step 1** | **Step 2** | **Step 3** | **Step 4** | **Step 5** | **Step 6** | **Step 7** | **Step 8** | **Step 9** |
| **FIB current** | 65 nA | 21 nA | 9.3 nA | 2.5 nA | 0.79 nA | 0.43 nA | 80 pA | 40 pA | 40 pA |
| **Lamella thickness**[&] | 60 µm | 30 µm | 20 µm | 10 µm | 5 µm | 2.5 µm | 1 µm | 500 nm | 100–150 nm |
| **Pattern X size** | 80 µm | 50 µm | 40 µm | 30 µm | 25 µm | 23 µm | 20µm | 20 µm | 20 µm |
| **Typical time** | 50 min | 30 min | 20 min | 60 min | 60 min | 30 min | 50 min | 30 min | 60 min |

[&] Lamellae thicknesses are measured from the front surface.

**Supplementary Table 2. Typical milling settings and milling times with CSEI-based locating prior to milling.** One surface of the final lamella was milled for applying CSEI-based locating prior to the milling steps described in the table. The milling described in the table was performed on a separate surface of the final lamella. All data were based on samples of ~50 μm thickness.

| | **locating** | **Coarse milling** | | | | | | **Fine milling** | |
|---|---|---|---|---|---|---|---|---|---|
| | **Step 1** | **Step 2** | **Step 3** | **Step 4** | **Step 5** | **Step 6** | **Step 7** | **Step 8** | **Step 9** |
| **FIB current** | 5 nA | 9.3 nA | 2.5 nA | 0.79 nA | 0.43 nA | 80 pA | 40 pA | 40 pA | 40 pA |
| **Lamella thickness**[&] | * μm | 20 μm | 10 μm | 5 μm | 2.5 μm | 1 μm | 1 μm | 500 nm | 100–150 nm |
| **Pattern X size** | 80 μm | 40 μm | 30 μm | 25 μm | 23 μm | 20 μm | 20μm | 20 μm | 20 μm |
| **Typical time** | * min | 60 min | 40 min | 40 min | 30 min | 50 min | 10 min | 30 min | 60 min |

* The time and cryo-lamellae thickness for the CSEI localization process are not fixed.

※ In this step, the rear of the lamellae is milled out at a 48 °angle.

**Supplementary Table 3. List of several typical examples.** Data on milled lamella size and number of tilt series collected on each lamella are shown for several samples with different initial thicknesses.

| Lamella no.[&] | Sample thickness[#] | Lamella area | Number of tilt series collected* |
|---|---|---|---|
| Lamella 1 | 60 μm | 20 μm×53 μm | 24 |
| Lamella 2 | 35 μm | 20μm×54 μm | 29 |
| Lamella 3 | 30 μm | 20 μm×60 μm | 22 |
| Lamella 4 | 52 μm | 20 μm×57 μm | 14 |

[&] These data represent only a subset of all cryoFIB experiments performed.

[#] Sample thickness (T) was based on the cross-sectional height (H) of the liver tissue after milling at an incidence angle of 48 °(T=H ×sin48 °/sin52 °).

* The statistical number of tilt series collected was not the maximum number of collections possible due to ice contamination issues and because only the area of interest was collected.
