## Supplementary Protocol for "CryoFIB milling large tissue samples for cryo-electron tomography"

This protocol is designed for preparing thin lamella starting from a thick, bulky sample. Mouse liver tissue will be used as an example in this protocol.

**Procedure**

**A. Initial sample preparation and fixation**

**1.** Dissect the liver from a mouse.

**2.** Wash the liver with phosphate buttered saline (PBS) pre-cooled to 4℃ or other optimal solution.

**3.** Cut the liver into small cuboids (with an edge length of 1-3 mm) with a razor blade.

**4.** **[Optional]** Some tissues tend to undergo rapid autolysis or cannot be frozen immediately after isolation from the organism. In these cases, chemical fixation is often performed using fixatives, such as paraformaldehyde (PFA) or glutaraldehyde (GA). The optimal fixative type and fixation time can vary depending on the sample. The dissected mouse liver tissue was fixed with 2.5% glutaraldehyde for 30 min at room temperature. Wash the tissue twice with pre-cooled PBS or other optimal solution after fixation.

**B. Pre-sectioning**

**1.** Pre-section the tissue using a vibratome or manual slicing. The recommended target thickness is 20–80 μm.

*__Note:__ The thickness can be adjusted according to experimental needs. Thinner samples require less cryoFIB milling time but may contain fewer target objects.*

**For vibratome sectioning (Leica VT1200S, Leica Microsystems):**

**a.** Heat 4% agarose until it melts.

**b.** Drop liquid agarose (approximately 1 mL) into a dish.

**c.** Place a tissue cuboid into the agarose and wait for solidification.

**d.** Mount the agarose-embedded sample onto the sample stage of the vibratome.

**e.** Fill the buffer tray of the vibratome with cold PBS or other optimal solution.

**f.** Set the vibratome parameters to 85 Hz sectioning frequency, 1 mm amplitude, and 0.5 mm/s sectioning speed.

*__Note:__ Vibratome parameters need to be optimized for different tissues.*

**g.** Set the vibratome thickness to 10–50 μm according to experimental needs.

**h.** Transfer the liver slices to cold PBS with a brush.

**For manual slicing:** Cut the tissue with a razor by hand. Since the thickness of manually sliced tissue may be thicker than expected, it is recommended to cut it as thinly as possible.

*__Note:__ To ensure tissue stability, it is important to keep pre-sectioning times short. Slicing with a vibratome typically takes more than 10 min. For tissues that are prone to autolysis, manual cutting is recommended.*

**C. Vitrification by high-pressure freezing**

**1.** Cool the high-pressure freezer (Leica EM HPM100, Leica Microsystems) to cryogenic temperature.

2. Ensure the freezing rate is above 12,000 K/s and the pressure is stable at approximately 300–350 ms.
3. Cool the cryo-ultramicrotome (Leica EM UC7+FC7, Leica Microsystems) from room temperature to -150 ℃.
4. Glow discharge the EM grids with parallel bars (Cu, 150 mesh, Zhongjingkeyi Technology, AG150P) using a glow discharger (PELCO easiGlow™, Glow Discharge Cleaning System). The parameters are set as follows: 0.40 mBar air pressure, 15 mA current, and 25 s glow discharge time. Grids with a small mesh number are recommended; 150-mesh grids have been used in the present work. Grids with or without lacey carbon film were tested, and both worked well.
5. Fill the dewar of the high-pressure freezer with clean liquid nitrogen.
6. Use a Perfect Loop (Diatome, DZ8) to transfer tissue slices to the cryoEM grids. The perfect loop is routinely used to pick up sections obtained by ultramicrotomy. Tweezers are not recommended in this step as they often damage the tissue slices.

   ***Note**: It is crucial to verify that the tissue slice lays flat on the grid and is well-adhered to the grid. If the tissue is not frozen solidly on the grid, it may fall off during subsequent operations.*
7. Fill the aluminum carrier (6 mm aluminum specimen carrier, Type A, 200 μm recess, Leica Microsystems) with 2-methylpentane.

   ***Note:** The cryo-protectant is not limited to 2-methylpentane. Other cryo-protectants can also be considered in different cases.*
8. Blot the grid using filter paper from the side without the sample to remove the excess buffer. Remove the Perfect Loop from the grid. Then, immediately place the grid into the aluminum carrier.

   ***Note:** To guarantee ideal vitrification, the excess buffer on the grid must be removed.*
9. Place a sapphire disc on the top of the aluminum carrier and immediately transfer the sandwiched assembly to the high-pressure freezer and vitrify the sample.
10. Transfer the sandwiched assembly to the cryo-ultramicrotome chamber previously cooled to -150 ℃ after high-pressure freezing. Note that the melting point of 2-methylpentane melts is -154 ℃.
11. Separate the grids from the aluminum carriers.
12. Keep the grids in the chamber for approximately 20 min to remove excess 2-methylpentane.
13. Transfer the grids containing tissue slices into grid boxes and store them in liquid nitrogen.

    ***Note:** Keep the liquid nitrogen clean and dry in the dewar of the high-pressure freezer, as this is essential for reducing ice contamination. The grids should not be placed in the cryo-ultramicrotome chamber for very long to avoid the accumulation of ice contamination on the tissue sample.*

**D. Preparation for cryoFIB milling**

***Note:** The cryoFIB milling procedure should be compatible with any cryoFIB* *instrument. The FEI Helios NanoLab G3 UC 992256 with Quorum cryo-system is used as an example in this protocol.*

1. Use red and black markers to mark the top (i.e. flat side), bottom, and outer side of the C-clip ring. The twelve o'clock direction is marked with red, and the three and nine o'clock directions with black (**Supplementary Protocol Fig. 1a and b**).
2. Cool the marked C-clip ring with dry liquid nitrogen.
3. Place the C-clip ring flat side down. Put the frozen grids containing tissue slices into the C-clip ring with the sample side down (**Supplementary Protocol Fig. 1a and b**).
4. Adjust the direction of the grid to ensure that the parallel grid bars are aligned to the red mark (**Supplementary Protocol Fig. 1a and b**) and assemble the Autogrid comprising the C-clip ring, grid, and C-clip following the standard procedure.
5. Mount the Autogrid onto a custom-made sample shuttle flat side up and adjust the direction of the Autogrid to make sure that the red marker points to the twelve o'clock direction (incident direction of the ion beam) (**Supplementary Protocol Fig. 1c**).

   ***Note:*** *The Autogrids must be mounted flat side up to reduce sheltering of the C-clip ring when milling from a shallow angle. If the direction of the parallel grid bars slightly deviates from the red mark, make sure that the parallel grid bars point to the twelve o'clock direction instead. This is a critical step in following the cryoET workflow.*
6. Transfer the shuttle to the prep-stage, which has been pre-cooled to -180°C in the prep chamber of the cryo-transfer system.
7. Elevate the temperature of the prep chamber to -150°C for 30–90 min to make sure any remaining 2-methylpentan on the grid is fully sublimated.
8. Set the prep chamber temperature to -180°C.
9. Transfer the shuttle into the SEM chamber for coating with the organometallic Pt protective layer.
   - **a.** Set the gas injection system (GIS) to 42°C.
   - **b.** Elevate the sample stage to a position such that the grid is 8 mm from the SEM tip.
   - **c.** Set the SEM parameters: acceleration voltage of 2 kV, 0.4 nA electron beam current, and a low magnification of 100x.
   - **d.** Insert the GIS needle and perform electron-beam-induced coating for 40 s with a continuous electron beam scanning over the entire coated area.
10. To improve the conductivity, transfer the shuttle to the prep chamber for sputter coating (5 mA, 60 s) to cover the organometallic Pt layer with a thin conductive Pt layer.
11. Transfer the shuttle back to the SEM chamber.

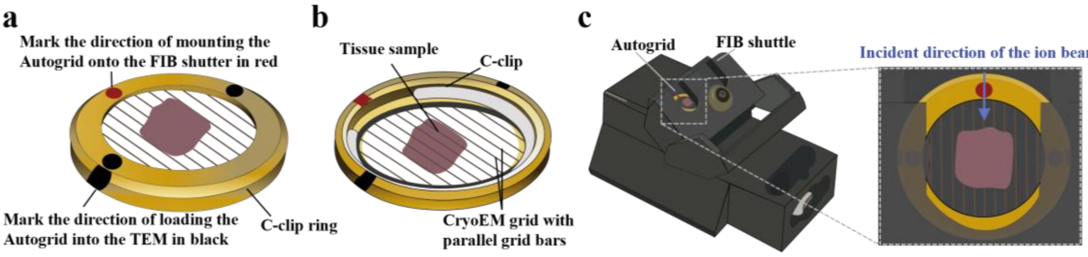


**Supplementary Protocol Figure 1. Schematic diagrams of the Autogrid prepared for cryoFIB. a** and **b,** An Autogrid composed of a C-clip ring, grid, and C-clip viewed from the top (**a**) and bottom (**b**). The top, bottom, and outer sides of the C-clip ring are marked in red and black. The red and black marks enable the correct orientation of the Autogrid in the FIB shuttle and Krios Cassette, respectively. The parallel grid bars are aligned to the red mark and the tissue on the cryoEM grid is facing up towards the flat top side of the Autogrid. **C.** The FIB shuttle loaded with an Autogrid. Magnified view showing a properly oriented Autogrid loaded into the custom-made sample shuttle. To allow for subsequent milling, the Autogrid is mounted onto the shuttle with the red mark oriented upwards (incident direction of the ion beam) and the tissue side facing outwards.

### E. Coarse milling

**1.** Tilt the sample stage such that the incident direction of the ion beam and the grid plane form a shallow angle (typically 18 °or less).

**2.** Acquire a low magnification SEM image of the whole grid and search for target positions.

*Note: Multiple lamellae can be milled together on the same grid. However, the selected regions on the grid generally need to comply with the following conditions: (1) the surrounding sample regions are firmly frozen, and are not warped or contain large cracks; (2) the region is between two grid bars to avoid the grid bars obstructing the ion and electron beam during cryoFIB milling and subsequent data collection; (3) the region has a clean surface (e.g., no ice contamination) to avoid the severe curtaining issue or the lamella being broken; and (4) multiple lamellae are separated as far as possible to maintain enough mechanical strength in the region adjacent to the lamella.*

**First sub-step of coarse milling:**

**3.** Tilt the sample stage so that the incident direction of the ion beam and the grid plane form a high angle of 48 °.

*Note: The milling angle in this step significantly influences the length of the lamellae produced (Fig. 3a). Milling at a high angle is particularly important for thick tissue samples.*

**4.** Acquire both FIB and SEM images of the target position.

**5.** Draw two windows of 80×100μm (width × height) above and beneath the target position under the FIB view using the Rectangle Pattern (named in the FIB user interface). The distance between the windows is set to tens of micrometers, typically 60–80 μm.

*Note:* *The size of the window is adjusted according to the thickness of the sample. The thicker the sample, the larger the window size should be, in order to avoid having the remaining bulky sample shelter the electron beam during cryoET tilt series acquisition. Generally, rectangular patterns of 80×100-150 μm are suggested for samples tens of micrometers thick.*

**6.** Set the ion beam current to 65 nA. The two windows are simultaneously milled. After the milling process, confirm that the volume at the milling positions is completely removed under FIB and SEM views. If not, repeat this operation.

***Note:*** *When the milled surface is too rough, a weaker beam (e.g., 9.3 nA) can be used to polish the surface by removing a thin layer. This surface will be the rear surface of the lamella. The rough surface may result in a discontinuous conductive Pt layer after the secondary sputter coating, which influences the function of the grounding wire.*

**Second sub-step of coarse milling:**

**7.** Tilt the sample stage so that the incident direction of the ion beam and the grid plane form a shallow angle (typically 18 °or less).

**8.** Perform lamella thinning by stepwise reduction of the ion beam current (typically from 21 nA, 9.3 nA to 2.5 nA) and moving the rectangular milling windows closer to the lamella. Gradually reduce the width of the two rectangular milling windows above and beneath the target position (under the FIB view) to create a stepped edge around the lamella. The width of the windows and the number of steps can be adjusted according to the thickness of the sample (see **Supplementary Table 1** for typical parameters).

***Note1:*** *In the case that the first sub-step is performed, once the sample is tilted back to 18 °or less, a fresh surface will be observed under the FIB view. Because the fresh surface is produced by FIB milling and not covered by an organometallic Pt layer, which is essential for protecting the lamella from damage during milling, the target position for further milling (i.e. the front surface of the final lamella) must not be chosen on the fresh surface (**Supplementary Protocol Fig.** 2).*

**9.** Disconnect one side of the lamella from the bulky sample by milling a window of 10 μm in width on either the right or the left side of the lamella.

***Note 1:*** *The vitrified bulky samples usually have strong inner stress. Therefore, we recommend disconnecting the lamella when the lamella is still thick (usually thicker than 15 μm) to avoid breakage during the milling process. In addition, the parallel grid bar can be easily bent, rendering the surrounding sample regions of the lamella unstable. Therefore, it is useful to mill a large gap (usually exceeding 10 μm in width) between the lamella and the remaining bulky sample in order to avoid crashing between them during future sample transfer.*

***Note 2:*** *Milling multiple lamellae (2–4) within a day is possible; however, when milling spans a day, contamination accumulation becomes a problem. To solve this problem, we recommend completing steps E1–E9 on all lamellae before proceeding to the subsequent milling steps.*

**10.** Draw three rectangular windows around the lamella. Mill by stepwise reduction of the ion beam current (typically 0.79 nA, 0.43 nA, and 80 pA) and moving the rectangular milling patterns closer to the target position.

**11.** Gradually reduce the width of the two rectangular windows above and beneath the target position (under the FIB view) to further create the stepped edge around the lamella. The width of the patterns and number of steps can be adjusted according to the thickness of the sample (see **Supplementary Table 1** for typical parameters).

**12.** Stop the milling procedure when the lamella is thinned to 1 μm thick.

***Note:*** *The width of the lamella (typical 10–20 μm) can be adjusted according to experimental needs.*

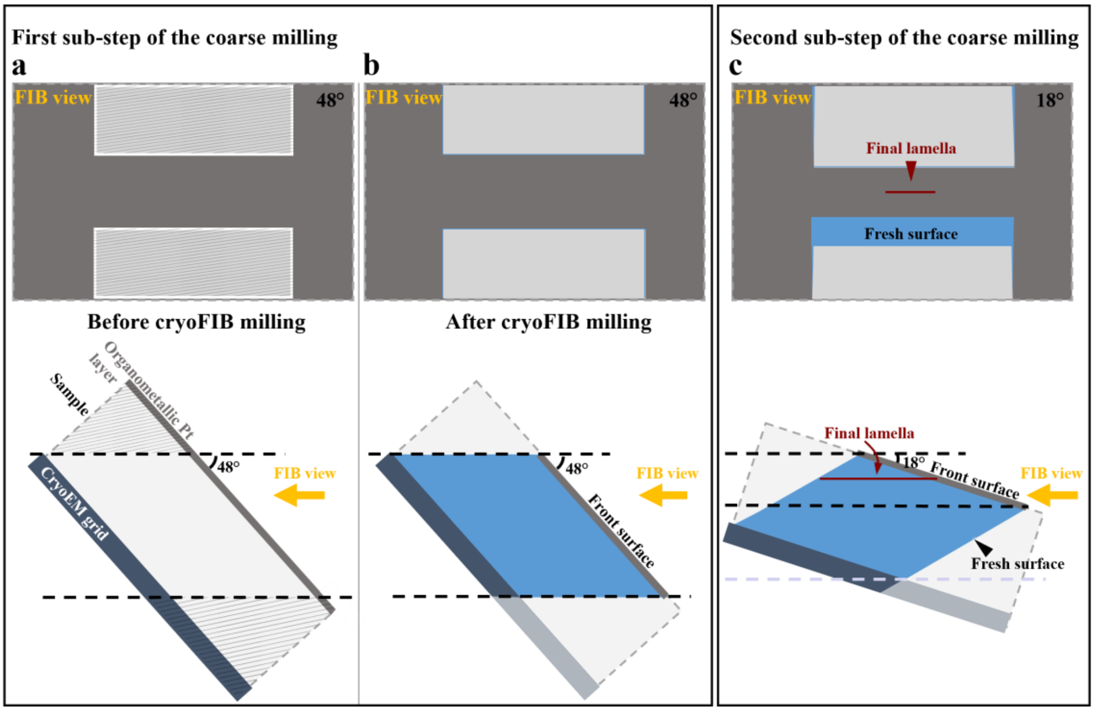


**Supplementary Protocol Figure 2. Schematic diagrams of the lamella during the coarse milling procedure.** The lamella is colored in blue, the bulky sample volume is represented as gray background, and the organometallic Pt layer is colored in dark gray. The conductive Pt layer is not shown in the diagrams. **a**–**b**, Schematic diagrams of the first sub-step of the coarse milling procedure viewed from FIB view (top) and side view (bottom). **a,** The sample is tilted such that the ion beam can mill along the incident angle of 48°. Two windows (white rectangles) were planned under the FIB view for the first sub-step of coarse milling. **b**, The two windows were milled at an incident ion beam angle of 48°, as specified in (**a**). **c,** Schematic diagrams of the second sub-step of the coarse milling procedure after (**b**) viewed from FIB view (top) and side view (bottom). The sample is tilted such that the ion beam can mill along the incident angle of 18°. A fresh surface is observed under FIB view without an organometallic Pt layer. The front surface of the final lamella is the top surface of the original sample covered by an organometallic Pt layer. The final lamella is represented by a red line.

**[Optional] CSEI locating procedure:**

**13.** When the objects of interest are sparsely distributed in a large bulky sample, we recommend locating the targets in the tissue sample using CSEI.

***Note:*** *CSEI can be used throughout the milling process.*

**a.** Randomly select a region (refer to step E2) and draw a window of 80 μm in width under cryoFIB view with a shallow beam angle (18 ° or less). The height can be adjusted according to the actual requirement.

***Note:*** *A large width means more structures can be observed but will require a longer milling time. Therefore, the width can be adjusted according to the actual requirement.*

**b.** Mill with proper ion beam current and continuously monitor the evolution of the milled surface by CSEI.

***Note:*** *A higher ion beam current improves the milling efficiency, thereby increasing the search volume, but causes more severe beam-induced damage. Once the object of interest is found on the milled surface, no further coarse milling should be applied at this surface to avoid removing the object of interest; this means that the damaged area caused by a high ion beam current cannot be removed as in steps E8-11. Therefore, the ion beam current should be optimized for each specific sample type to properly balance the quality of the lamella and the milling efficiency. It is highly recommended to use a lower ion beam current (e.g., ~5 nA or less) throughout the CSEI process to minimize beam-induced damage. If the object of interest is large enough, it is possible to coarsely locate the object using a higher ion beam current and then polish the surface using a lower ion beam current.*

**c.** Repeat steps **a**–**b**. The milling process is complete as soon as the object of interest is located.

***Note:*** *Multiple regions can be simultaneously or sequentially searched using the CSEI procedure.*

**d.** Mill on the opposite side of the CSEI by stepwise reduction of the ion beam current (typically from 9.3 nA, 2.5 nA, 0.79 nA to 80 pA) and moving the rectangular milling window closer to the lamella. The width of the rectangular milling window should be gradually reduced to create a stepped edge on one side of the lamella (see **Supplementary Table 2** for typical parameters).

***Note:*** *The highest ion beam current is usually reduced to 9.3 nA or lower in this step to avoid beam-induced damage.*

**e.** Disconnect one side of the lamella from the bulky sample by milling a window of 10 μm in width on either the right or the left side of the lamella.

***Note:*** *The milling procedure is similar to steps E8–11.*

**f.** Stop the milling procedure when the lamella is thinned to 1 μm thick.

**g.** When using the CSEI procedure ahead of coarse milling, the rear surface of the lamella cannot be coated with the metallic Pt layer during the second sputter coating process (**Supplementary Fig. 6b**). Therefore, a sloped surface that can be effectively coated by sputter coating should be created as follows:

**i.** Tilt the sample stage so that the incident direction of the ion beam and the grid plane form a high angle of 48°.

**ii.** Draw a rectangular milling window (20×5 μm) at the rear end of the lamella and mill using a very low ion beam current (24–40 pA).

***Note:*** *The width of the window depends on the lamella width. The top surface of the lamella, which is not protected by the organometallic Pt layer, will be exposed to the ion beam when tilted to 48°. This surface can be damaged by the ion beam during FIB imaging and milling. Therefore, try to minimize unnecessary FIB imaging and lower the ion beam current for milling.*

**iii.** Tilt the sample stage back to the previous position and tilt angle. Check that the rear surface of the lamella is removed under SEM view.

***Note:*** *When CSEI is used ahead of coarse milling to locate targets, the first sub-step of coarse milling (steps E3–E6) should be skipped to avoid beam-induced damage by the strong ion beam. In such cases, we recommend preparing thin samples, typically not thicker than 30 μm, during the previous pre-sectioning step to avoid long milling times.*

**Final sputter coating:**

**14.** Transfer the shuttle to the prep chamber for sputter coating (5 mA, 60 s).

***Note:*** *The coating should cover the front, top, and rear surfaces of the lamella; this is critical for eliminating charge accumulation during data collection.*

**F. Fine milling the lamella**

**1.** Transfer the shuttle in the prep chamber back to the SEM chamber and adjust the grid back to the previous position and tilt angle.

**2.** Reduce the ion beam current to 40 pA. Use the Cleaning Cross-Section Pattern (named in the FIB user interface) in all the following milling operations to relieve curtaining on the lamella.

**3.** Draw a window of 20 μm × ~500 nm on the bottom side of the lamella, then perform the milling. The milling direction of the window is set from bottom to top.

**4.** Draw four parallel windows of 3 μm × ~100 nm on the top side of the lamella, then perform the milling. The distance between adjacent windows is 2 μm. The milling direction is set from bottom to top. Confirm that four notches (i.e. the furrows-to-be) can be clearly observed under the FIB view. If not, repeat this operation.

***Note:*** *The width of the milling windows can be adjusted according to the illumination area for data collection. This procedure is performed to mark the milling positions for the furrows.*

**5.** Increase the magnification of FIB imaging such that one notch fills the field of view. Draw two windows of 3 μm in width on the top and bottom sides of the lamella at the position of the notch, then perform the milling to make one furrow.

***Note:*** *The thickness of the furrow can be measured more precisely at higher* *magnification.*

6. **6.** Continuously monitor the milling area by live SEM imaging to make sure the milling is proceeding as expected. Pause the milling and refresh the FIB image to measure the thickness of the furrow frequently.

7. **7.** Stop the milling after reaching the target thickness.

   ***Note:*** *The organometallic Pt layer on the front surface of the lamella can be clearly viewed by SEM imaging. This Pt layer will form a meniscus at the front of the furrow (**Fig. 4d**) and continuously narrow when milling the furrow. The milling should be stopped immediately if a notch is observed on the meniscus to avoid breaking the furrow.*

8. **8.** Repeat steps F5–7 to make four furrows. The furrows should be milled in order beginning from the disconnected end to the fixed end of the lamella.

**H. Tilt series data collection**

1. **1.** Load the Autogrid with milled samples into a cryo-electron microscope. The black mark on the outer side of the Autogrid is oriented upward in the Krios Cassette, ensuring that the milling direction of lamellae is perpendicular to the tilt axis for cryoET data collection.

2. **2.** Select a position of interest in one of the four furrows for data collection. The tracking/focusing area can be set in a furrow adjacent to the position of interest.

   ***Note:*** *The electron beam should always touch a ridge during data-acquisition illumination to reduce charging.*
